# Elasticity and Thermal Stability are Key Determinants of Hearing Rescue by Mini-Protocadherin-15 Proteins

**DOI:** 10.1101/2024.06.16.599132

**Authors:** Pedro De-la-Torre, Haosheng Wen, Joseph Brower, Karina Martínez-Pérez, Yoshie Narui, Frank Yeh, Evan Hale, Maryna V. Ivanchenko, David P. Corey, Marcos Sotomayor, Artur A. Indzhykulian

**Affiliations:** Department of Otolaryngology - Head and Neck Surgery, Harvard Medical School and Massachusetts Eye and Ear, 243 Charles St, Boston, MA, USA; Department of Chemistry and Biochemistry, The Ohio State University, 484 W. 12th Avenue, Columbus, OH, USA; Biophysics Program, The Ohio State University, 484 W. 12th Avenue, Columbus, OH, USA; Biology Program, Department of Basic Sciences, Universidad del Atlántico, Cra 30 # 8-49, Puerto Colombia, 081007, Colombia; Center for Electron Microscopy and Analysis, The Ohio State University, 1275-1305 Kinnear Road, Columbus, OH, USA; Department of Neurobiology, Harvard Medical School, 200 Longwood Ave, Boston, MA, USA; Speech and Hearing Biosciences and Technology graduate program, Harvard University, Cambridge, MA, USA

## Abstract

Protocadherin-15 is a core protein component of inner-ear hair-cell tip links pulling on transduction channels essential for hearing and balance. Protocadherin-15 defects can result in non-syndromic deafness or Usher syndrome type 1F (USH1F) with hearing loss, balance deficits, and progressive blindness. Three rationally engineered shortened versions of protocadherin-15 (mini-PCDH15s) amenable for gene therapy have been used to rescue function in USH1F mouse models. Two can successfully or partially rescue hearing, while another one fails. Here we show that despite varying levels of hearing rescue, all three mini-PCDH15 versions can rescue hair-cell mechanotransduction. Negative-stain electron microscopy shows that all three versions form dimers like the wild-type protein, while crystal structures of some engineered fragments show that these can properly fold and bind calcium ions essential for function. In contrast, simulations predict distinct elasticities and nano differential scanning fluorimetry shows differences in melting temperature measurements. Our data suggest that elasticity and thermal stability are key determinants of sustained hearing rescue by mini-PCDH15s.

## INTRODUCTION

The first step in auditory perception is the conversion of a sound stimulus into an electrical signal by hair cells of the inner ear, which are named for their mechanosensitive bundle of actin-based microvillus-like projections called stereocilia. Long protein filaments known as “tip links”^1–5^ interconnect stereocilia and are essential for hair-cell function (**Fig. 1a)**. Mechanical forces from sound or acceleration due to head movements and gravity lead to stereocilia bundle displacement and tensioning of tip links that pull on mechanosensitive ion channels enabling hair-cell mechanoelectrical transduction function^6–8^. Tip links are essential for mechanotransduction, they can be regenerated after chemical disruption^9,10^, and their function is calcium (Ca^2+^)-dependent.

**Fig. 1.**
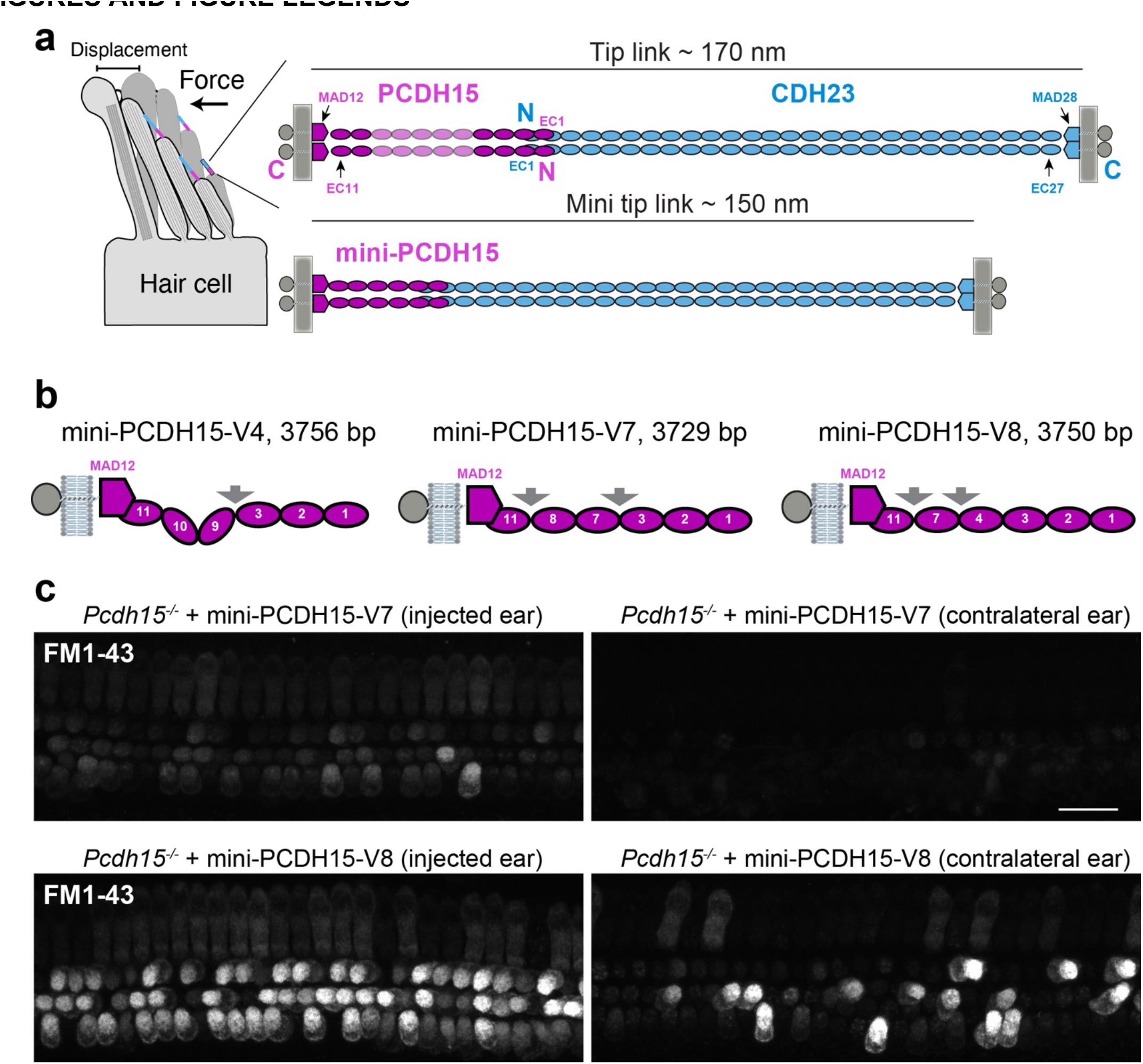
**| Mini-PCDH15-V7 and -V8 rescue hair-cell mechanotransduction function in *Pcdh15^-/-^* mice. a**, Hair-cell stereocilia bundle and location of tip links. Force induces deflection of stereocilia bundles and stretches tip links initiating the mechanotransduction process. CDH23 (blue) and PCDH15 (magenta) interact tip-to-tip to form the tetrameric tip-link complex. Mini-tip-links designed via deletion of EC repeats from PCDH15 form mini-PCDH15s. **b,** Schematics of mini-PCDH15 proteins highlighting engineered EC-EC linkers (arrows). Mini-gene sizes of each ectodomain are indicated in base pairs (bp). **c**, Cochlear hair cells of *Pcdh15^-/-^* mice treated at P1 with inner ear delivery of AAV-mini-PCDH15-V7 or - V8 show restored mechanotransduction function as assessed by application of FM1-43 dye loading 4 days post-injection. Some hair cells from the contralateral ear also report FM1-43 uptake, likely due to the AAV crossing over to the other ear. *Scale bar*, 20 µm.

Tip links are formed by two pairs of large Ca^2+^-binding unconventional cadherins, cadherin-23 (CDH23), with its extracellular domain forming the upper ∼2/3 of the tip link^1,2,11^ and protocadherin-15 (PCDH15) forming the lower

∼1/3^12^. PCDH15 is also detected in the retina and brain^13^, although its overall function beyond inner-ear mechanotransduction is less understood^14–17^. Defects or absence of the PCDH15 protein can cause non-syndromic deafness DFNB23 or Usher syndrome type 1F (USH1F), a disease characterized by congenital hearing loss, balance deficit, and progressive blindness^16,18–21^. Several reports have also implicated PCDH15 in brain function^22–26^ and disorders including autism, bipolar disorders, and schizophrenia^27–30^.

Given the link between *PCDH15* mutations and various sensory disorders, gene therapy technologies to deliver a functional copy of the *PCDH15* coding sequence (> 6 kb) have been explored. Adeno-associated viruses (AAV) have been shown to be efficient and effective for gene therapy^31–35^, but these have an inadequate packaging capacity (< 4.7 kb^36^) to carry the full length *PCDH15* coding sequence in a single AAV. Dual-AAV approaches have been successfully implemented in several gene therapy studies and clinical trials^37–43^, including two aiming to express full-length PCDH15^44,45^. However, shortening PCDH15 by removing possibly unnecessary domains could enable a more efficient transgene delivery to patients by a single AAV.

PCDH15 has 11 extracellular cadherin (EC) repeats interconnected by Ca^2+^-binding acidic residues that stabilize the protein^46^. The N-terminus of PCDH15 and CDH23 interact to form tip links, while parallel *cis* homodimerization of PCDH15 is mediated by the EC2-3 fragment and a membrane-adjacent domain (MAD12^47^, also called the SEA^48^, PICA^49^ or EL^50^ domain; **Fig. 1a)**. In a previous study^51^, we reported eight shortened PCDH15 proteins (referred to as mini-PCDH15s) lacking 3 to 5 EC repeats (∼300 to ∼550 amino acids) of the native mouse (*mm*) PCDH15 sequence (species will be assumed to be *mm* in the text below unless otherwise noted). All variants were rationally engineered to keep structural features important for the interaction of PCDH15 with CDH23, for PCDH15 parallel dimerization, and for Ca^2+^ binding. One of these variants, mini-PCDH15-V4, successfully rescued hearing function as measured by brainstem recording when introduced into the cochlea at postnatal day 1 (P1) using AAV delivery in three PCDH15-deficient mouse lines. Another variant, mini-PCDH15-V7, partially rescued hearing, while mini-PCDH15-V8 failed.

To understand the varying degrees of auditory function rescue by mini-PCDH15s, we first tested their ability to mediate mechanotransduction function in cochlear hair cells of *Pcdh15^-/-^* mice transduced with AAVs encoding mini-PCDH15-V4, -V7, or -V8. All three mini-PCDH15s rescued hair-cell mechanotransduction, consistent with our previous experiments^51^ showing that these variants interacted with CDH23 and suggesting that sustained rescue of hearing, or the lack of it, may depend on other specific structural and dynamical features of the mini-PCDH15s. Negative-stain electron microscopy (EM) also confirmed that all three variants form parallel dimers, albeit with some displaying more varied conformations than others. X-ray crystal structures of engineered EC-EC linkers showed that these can fold and can bind Ca^2+^ as expected, indicating that differences in structural details alone may not explain the variability in hearing rescue by mini-PCDH15s. In contrast, steered molecular dynamics (SMD) simulations showed distinct elasticity for the ectodomains of the three mini-PCDH15 variants, while nano differential scanning fluorimetry (nanoDSF) revealed differences in melting temperature measurements. These data suggest that tip-link elasticity and thermal stability are key determinants of sustained hearing rescue by mini-PCDH15s, paving the way to further improve AAV-based mini-gene therapies involving PCDH15 and other large cadherin proteins.

## RESULTS

### Three mini-PCDH15s with differential hearing-rescue abilities facilitate hair-cell mechanotransduction

In our previous study, auditory brainstem response (ABR) and distortion product otoacoustic emissions (DPOAE) recordings were used to assess the potential of the shortest mini-PCDH15 variants (-V4, -V7 and -V8; **Fig. 1b**) in rescuing auditory function in mice. These experiments showed varied levels of functional rescue^51^. To ascertain the potential for these three engineered mini-PCDH15 variants to mediate hair-cell mechanotransduction function, we injected neonatal *Pcdh15^-/-^* mice at postnatal day 1 (P1) with AAVs encoding mini-PCDH15-V7 or -V8 using round window membrane delivery method. Four days post-injection, the cochlear sensory epithelia were acutely dissected and briefly treated with a positively charged styryl FM1-43 dye known to enter hair cells through functional mechanotransduction channels^52–54^. While the hair cells of *Pcdh15^-/-^* mice do not take up FM1-43^51^ and are not mechanosensitive, AAV-mediated expression of mini-PCDH15-V7 and -V8 in injected ears resulted in restoration of FM1-43 uptake in most hair cells (**Fig. 1c**). This is similar to what we found for mini-PCDH15-V4 tested previously^51^. Thus, despite the varied level of hearing rescue in PCDH15- deficient mice expressing one of the three mini-PCDH15 variants, the ability of these mini-PCDH15s to mediate mechanotransduction function suggests an intricate interplay between their structure and ability to restore function, warranting further investigation.

### Negative stain electron microscopy shows that mini-PCDH15 ectodomains form *cis* dimers

Analyses of crystal structures^46^ as well as of negative stain EM^49^, cryo-EM^50^, and cryo-electron tomography (cryo-ET)^55^ data support parallel *cis* dimerization of the PCDH15 ectodomain *in vitro* and in hair cells *ex vivo*.

There are two ectodomain regions that interact to facilitate *cis* PCDH15-dimer formation: the EC2-3^46,49^ fragment near the N-terminus forming an “X” dimer and the EC11-MAD12^47,50^ fragment near the ectodomain C-terminus. Our previous size-exclussion chromatography (SEC) analyses suggested that dimerization of all mini-PCDH15 variants was preserved despite shortening, but provided little details on whether this dimerization involved the same parallel interfaces expected for the wild-type (WT) ectodomain.

To investigate the *cis* homodimerization of mini-PCDH15-V4, -V7, and -V8, we obtained negative-stain EM micrographs for each in the presence of 5 mM Ca^2+^. Class averages for mini-PCDH15-V4 showed the expected *cis-*homodimer architecture with two protomers in diverse conformations **(Fig. 2)**. We identified two lobes that likely correspond to the EC1-2 repeats forming an open-scissor conformation coupled to the EC3 repeat, similar to what has been reported for the structure of the PCDH15 EC1-3 fragment alone^49^ and for the tetrameric structure of PCDH15 EC1-3 in complex with CDH23 EC1-2^46^. At the opposite end we often found a bright electron-dense region that we have assigned to the EC11-MAD12 dimeric fragment **(Fig. 2)**. The EC2-3 dimeric point is characterized by an increased electron density area followed by a kink and a separation of the two protomers, likely at the noncanonical EC9-10 Ca^2+^-free linker region, which is absent in mini-PCDH15-V7 and - V8. Some class averages of mini-PCDH15-V4 showed conformations that we labeled “*L*” due to their apparent shape. The kinks associated to this conformation likely correspond to the engineered EC3-EC9 and the native EC9-10 regions **(Fig. 2**, and **Movie S1)**. It is likely that the engineered EC3-EC9 linker displays increased flexibility, perhaps due to decreased affinity for Ca^2+^. If so, this linker would provide increased elasticity to mini-PCDH15-V4, similar to the EC5-6 linker within the WT PCDH15 ectodomain^46^.

**Fig. 2.**
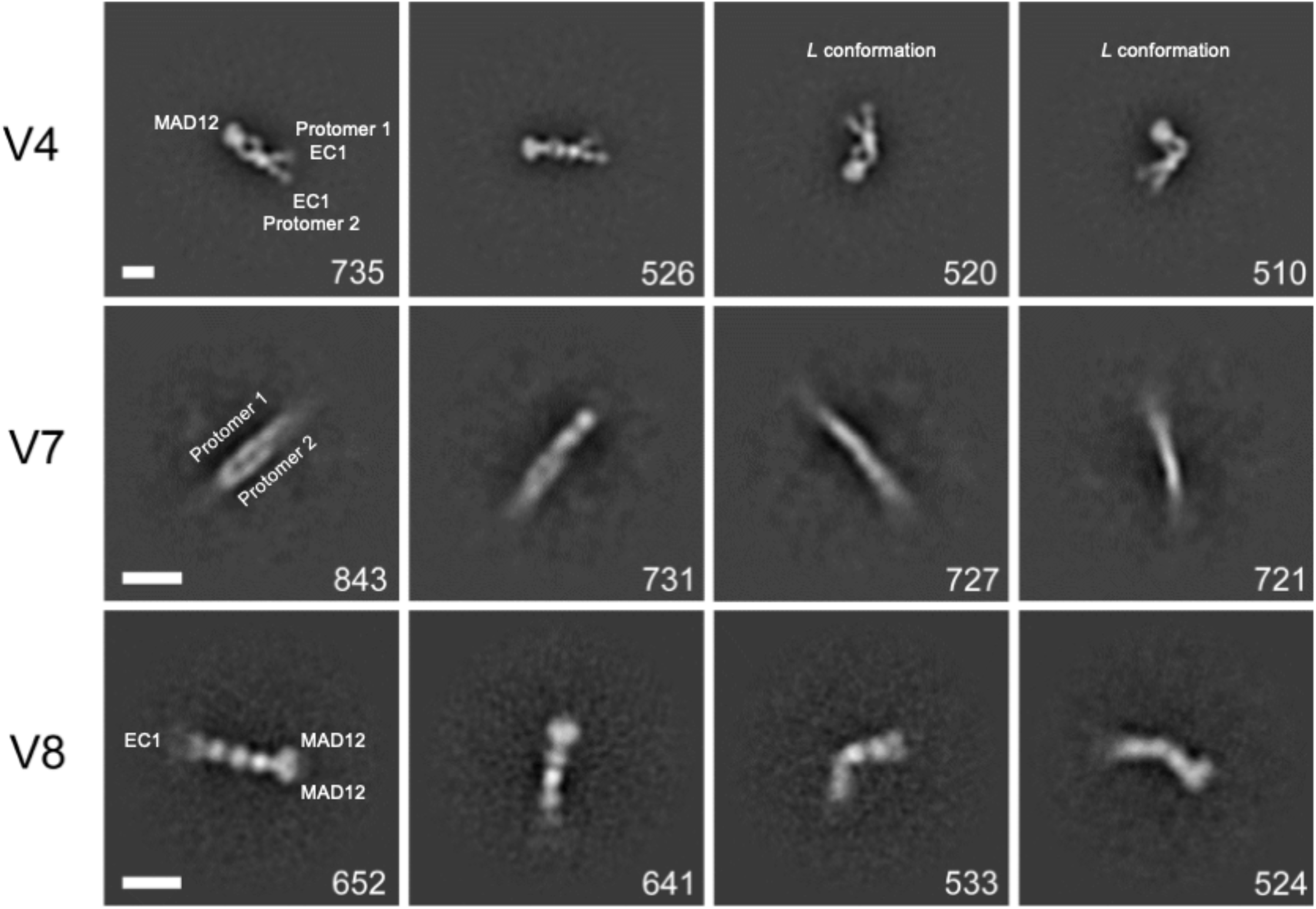
**| 2D Class averages of mini-PCDH15 from negative-stain EM**. Classification of ∼10,000 particles shown in order of decreasing abundance for mini-PCDH15-V4 (*top*), mini-PCDH15-V7, and mini-PCDH15-V8 in the presence of 5 mM Ca^2+^. All mini-PCDH15 versions appear to be dimeric. The number of individual images used for the 2D-class averages is shown at the lower right of each image. Box size is 944 Å x 944 Å (*top*); 700 Å x 700 Å (*middle and bottom*). Scale bar indicates 100 Å.

Class averages for mini-PCDH15-V7 and -V8 showed dimers as well **(Fig. 2)**. However, identification of EC repeats and location of N- and C-termini in the 2D class averages for mini-PCDH15-V7 was difficult due to a slight feathering of both ends of the dimer. In contrast, the 2D class averages of mini-PCDH15-V8 showed straight and bent conformations for the ectodomain. At its N-terminus, we found two attenuated lobes likely for EC1-2, followed by the engineered EC3-EC4-EC7-EC11-MAD12 protein segment where a bright electron dense region was assigned as the dimeric EC11-MAD12 **(Fig. 2)**. These 2D micrographs suggest that the engineered EC7-EC11 or EC8-EC11 linkers might be prone to bending.

In general, the 2D class averages confirm that mini-PCDH15-V4, -V7, and -V8 form *cis* homodimers. Importantly, we observed kinks for mini-PCDH15-V4, straight conformations for mini-PCDH15-V7, and some bent conformations for mini-PCDH15-V8.

### Crystal structures reveal that engineered EC-EC linkers fold and can bind Ca^2+^

Previous structural work characterized all parts of the WT PCDH15 ectodomain at high resolution, including all its EC-EC linker regions. Some of these regions are canonical and have three Ca^2+^-binding sites labeled 1, 2 and 3 (EC1-2, EC4-5, EC6-7, EC7-8, EC8-9), while others are non-canonical and bind two Ca^2+^ ions (EC2-3, EC3-4), one (EC5-6), or none (EC9-10). The mini-PCDH15 proteins feature rationally engineered linkers that are not found in nature, including EC3-EC9 for mini-PCDH15-V4, EC3-EC7 and EC8-EC11 for mini-PCDH15- V7, and EC4-EC7 and EC7-EC11 for mini-PCDH15-V8 **(Fig. 1b)**. To understand how these engineered linkers fold and bind Ca^2+^ ions we obtained X-ray crystal structures for EC3-EC7 and EC4-EC7. For all mini-PCDH15 proteins, residue numbering was preserved to be in agreement with the mature (no signal peptide) WT *mm* PCDH15 ectodomain^46^ (NCBI accession number NP_001136214.1, see *Methods*).

The structure for the mini-PCDH15 EC3-EC7 linker was solved and refined to 1.94 Å resolution **(Table 1)**. There was one protomer in the asymmetric unit comprising amino acids p.D238–370Q and p.A700–D799 from the EC3 and EC7 repeats, respectively **(Fig. 3)**. The nonconsecutive EC3 and EC7 repeats were interconnected with the noncanonical cadherin Ca^2+^-binding motif p.366DENNQ from the native EC3 C-terminal end followed by residue p.A700 from EC7. The EC3-EC7 structure adopts a bent conformation with a typical cadherin Greek-key fold featuring seven *β*-strands for each repeat **(Fig. 3)**. Root mean square deviation (RMSD) values for EC3 repeats from our structure and previous WT structures were < 1.1 Å **(Table S1)**. Similarly, RMSD values for EC7 repeats were < 0.65 Å **(Table S1)**. There is one sodium ion (Na^+^) bound at the N-terminus of EC3 in a solvent-exposed site 3 **(Fig. 3a**, *and inset*), while there are no ions at the C-terminal end region of the EC7 repeat. The EC3-EC7 linker region has one bound potassium ion (K^+^) at site 1 and one Ca^2+^ ion bound at site 3. No ions were observed at site 2 **(Fig. 3a**, *and inset*). The bent EC3-EC7 structure does not show the typical features expected for a canonical linker region but reveals proper folding for each EC repeat and a semi-canonical connection between them.

**Fig. 3.**
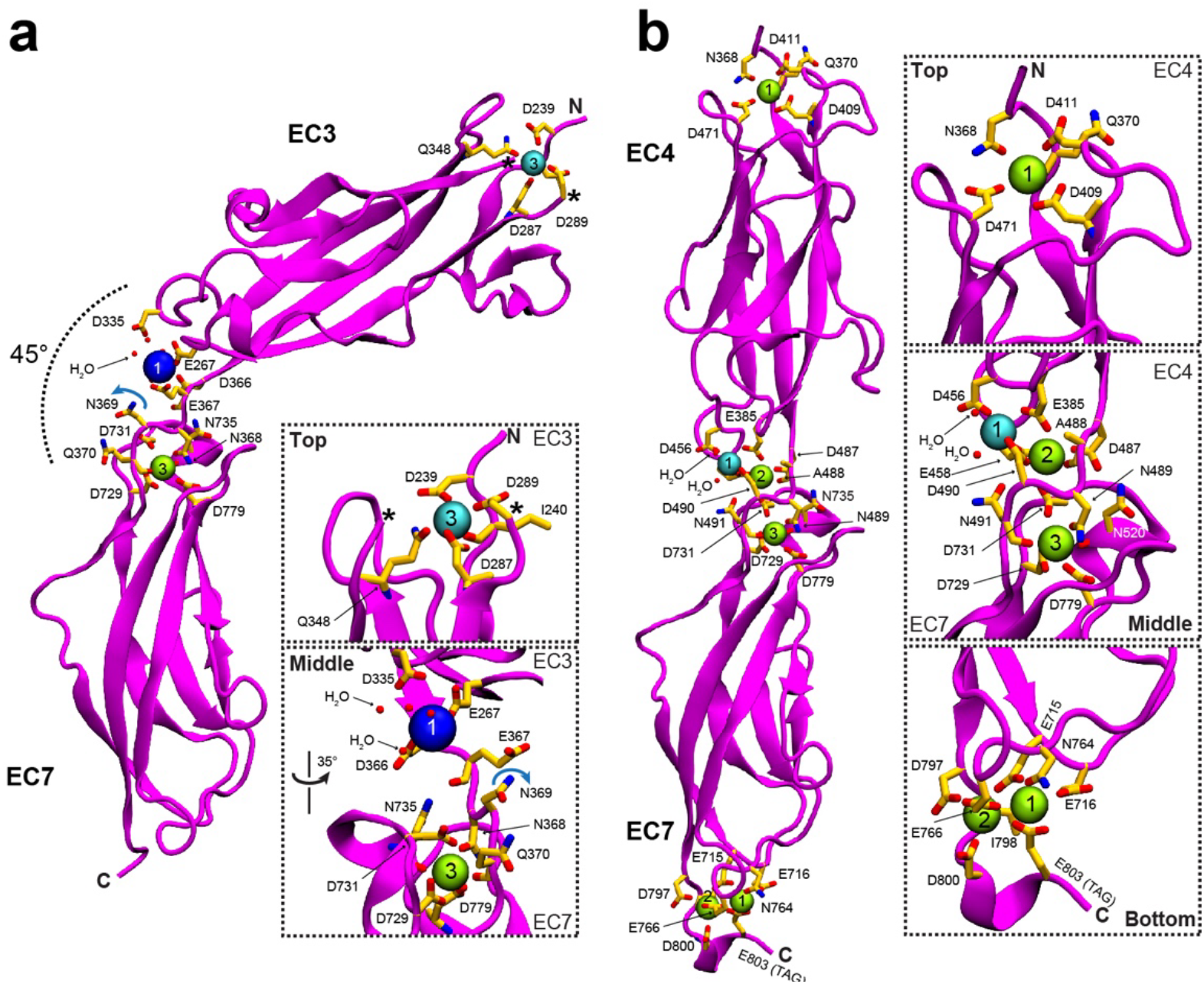
| Crystal structure of engineered EC-EC linkers. a-b, X-ray crystal structures of the engineered EC3-EC7 (mini- PCDH15-V7) and EC4-EC7 (chain A, mini-PCDH15-V8) fragments, respectively. Protein is shown in magenta. Ca^2+^, K^+^, and Na^+^ ions are shown as green, cyan, and blue spheres. Black star indicates a missing loop. Water molecules completing the coordination sphere are shown as small red spheres. Residues involved in ion coordination are shown as yellow sticks and labeled.

**Table 1.** | Data collection and refinement statistics

| **Data collection** | ***mm* PCDH15 EC3-EC7** | ***mm* PCDH15 EC4-EC7** |
| --- | --- | --- |
| Space group | *P* 3<sub>2</sub> 2 1 | *P* 1 2<sub>1</sub> 1 |
| Unit cell parameters |  |  |
| *a*, *b*, *c* (Å) | 84.60, 84.60, 76.89 | 49.276, 50.613, 88.263 |
| $\alpha$, $\beta$, $\gamma$ (°) | 90.0, 90.0, 120.0 | 90.0, 90.0, 94.01 |
| Molecules per asymmetric unit | 1 | 2 |
| Beam source | APS-24-ID-E | APS-24-ID-E |
| Date of data collection | 21-SEPT-2022 | 16-DEC-2022 |
| Wavelength (Å) | 0.97918 | 0.97918 |
| Resolution limit (Å) | 1.94 | 1.63 |
| Unique reflections | 23952 | 54156 |
| Completeness (%) | 99.9 (100) | 93.6 (96.2) |
| Redundancy | 11.6 (10.4) | 2.8 (2.9) |
| *I* / $\sigma(I)$ | 31.929 (1.86) | 10.956 (2.293) |
| *R*<sub>merge</sub> | 0.051 (0.852) | 0.074 (0.406) |
| *R*<sub>meas</sub> | 0.053 (0.897) | 0.091 (0.493) |
| *R*<sub>pim</sub> | 0.015 (0.277) | 0.052 (0.277) |
| *CC*<sub>1/2</sub> | 1.0 (0.728) | 0.997 (0.841) |
| *CC*<sup>*</sup> | 1.0 (0.918) | 0.999 (0.956) |
| **Refinement** |  |  |
| Resolution range (Å) | 50.0 – 1.94<br>(1.97 – 1.94) | 50.0 – 1.64<br>(1.67 – 1.64) |
| *R*<sub>work</sub> (%) | 18.40 (32.5) | 22.276 (29.0) |
| *R*<sub>free</sub> (%) | 22.30 (33.0) | 27.354 (37.0) |
| Residues (atoms) | 224 (1776) | 444 (3508) |
| Water molecules | 125 | 262 |
| R.M.S. deviations |  |  |
| Bond lengths (Å) | 0.0114 | 0.0106 |
| Bond angles (°) | 1.7127 | 1.7032 |
| *B*-factor average |  |  |
| Protein | 45.93 | 25.08 |
| Ligand/ion | 48.95 | 21.29 |
| Water | 46.39 | 28.92 |
| **Ramachandran Plot Region (PROCHECK)** |  |  |
| Most favored (%) | 94.2 | 91.8 |
| Additionally allowed (%) | 4.7 | 7.9 |
| Generously allowed (%) | 1.1 | 0.3 |
| Disallowed (%) | 0.0 | 0.0 |
| **PDB ID code** | **8TON** | **8UMZ** |

A structural comparison between the native non-canonical EC3-4 linker region (PDB: 5T4M), the native canonical EC6-7 linker region (PDB: 6BWN), and the engineered EC3-EC7 linker region reveals key differences that may explain ion occupancy at these linkers. In the p.366DENNQ linker motif of EC3-EC7, the asparagine p.N369 is pointing away from site 2 **(Fig. 3a**, *and inset*) while the same residue in the native EC3-4 linker (PDB: 5T4M) coordinates a Ca^2+^ ion at site 2^46^, and in canonical linkers with a DXNDN motif (X represents any amino acid) the equivalent aspartate coordinates Ca^2+^ ion at sites 2 and 3. Our results suggest that the EC3-EC7 structure is most similar to the native non-canonical EC3-4 than to the native canonical EC6-7 structures, which may explain its bent conformation and perhaps increased flexibility^46^.

The structure of mini-PCDH15 EC4-EC7 was solved at 1.63 Å resolution and showed two protomers in the asymmetric unit. Models include residues p.E367-N491 and p.A700–D800 (chain A), and p.E367-N491 and p.A700–N801 (chain B) from EC3 and EC7, respectively **(Fig. 3b)**. The nonconsecutive EC4 and EC7 repeats were interconnected with the canonical cadherin Ca^2+^-binding motif p.487DANDN from the native EC4 C-terminal end followed by residue p.A700 from EC7. The EC4-EC7 structure shows clear straight conformations for both protomers, similar to other native structures^46^. RMSD values for EC4 repeats from our structure and previous WT structures were < 0.93 Å **(Table S1)**. There is one Ca^2+^ ion bound at the N-terminal top of EC3 in a solvent-exposed site 3 **(Fig. 3b**, *and insets*) and two Ca^2+^ ions at the C-terminal end region of the EC7 repeat, assigned to sites 1 and 2 and interacting with residue p.E588 from the *XhoI* cloning site right before the 6X-Histidine tag. The EC4-EC7 linker region has one Na^+^ ion at position 1, and Ca^2+^ ions at sites 2 and 3 **(Fig. 3b)**. The straight EC4-EC7 structure shows typical features expected for a canonical linker region with proper folding for each EC repeat.

Overall, our structures indicate that engineered linker regions can be bent or straight and can have both noncanonical and canonical linker regions with EC repeats that adopt typical cadherin folds. Interestingly, AlphaFold2 (AF2) and AlphaFold3 (AF3) predictions for the EC3-EC7 and EC4-EC7 fragments are consistent with the architecture of the individual repeats (**Fig. S1, Table S1**, and **S4)** with some differences related to ion occupancy and degree of bending at the linker regions. Nevertheless, our structures validate the use of structural information from WT proteins as well as homology and machine learning techniques to engineer cadherin fragments where EC repeats are placed next to each other to form new EC-EC linker regions not found in nature.

### Equilibrium and SMD simulations predict differential elasticity for mini-PCDH15 ectodomains

Considering that all three mini-PCDH15s facilitate hair-cell mechanotransduction, form WT-like dimers and that engineered linker regions form canonical and noncanonical EC-EC connections as seen in WT PCDH15, we wondered if differential dynamics and elasticity of mini-PCDH15s could be the cause of their differential ability to rescue hearing. To study the dynamics and elasticity of mini-PCDH15 proteins, we built models of the full-length monomeric ectodomains of mini-PCDH15-V4, -V7, and -V8 and tested their elasticity *in silico*.

Because we have solved the atomic structures for two out of five engineered EC-EC linker regions and our structures of engineered EC-EC fragments were partially consistent with prior WT structures and with predictions from AlphaFold, we used two approaches to model the mini-PCDH15 ectodomains. In the first one, we used engineered linker regions predicted by AF2 within experimentally derived models of the rest of the ectodomain. In the second approach, engineered linker regions were modeled after aligning (AL) experimental crystal structures, and these were kept as parts of experimentally derived models of the rest of the ectodomain (see *Methods*). We reasoned that different starting models would lead to more sampling and that model-independent results would lead to robust predictions. These six mini-PCDH15 models were labeled as mini-PCDH15-V4 AF2, mini-PCDH15-V4 AL, mini-PCDH15-V7 AF2, mini-PCDH15-V7 AL, mini-PCDH15-V8 AF2, and mini-PCDH15-

V8 AL **(Fig. 4a, d, g, j, m, p)**. Details of Ca^2+^ placement and coordination are discussed in the Supplementary Materials. All models were solvated (150 mM KCl) and used to run 100-ns long all-atom equilibrium molecular dynamics (MD) simulations.

**Fig. 4.**
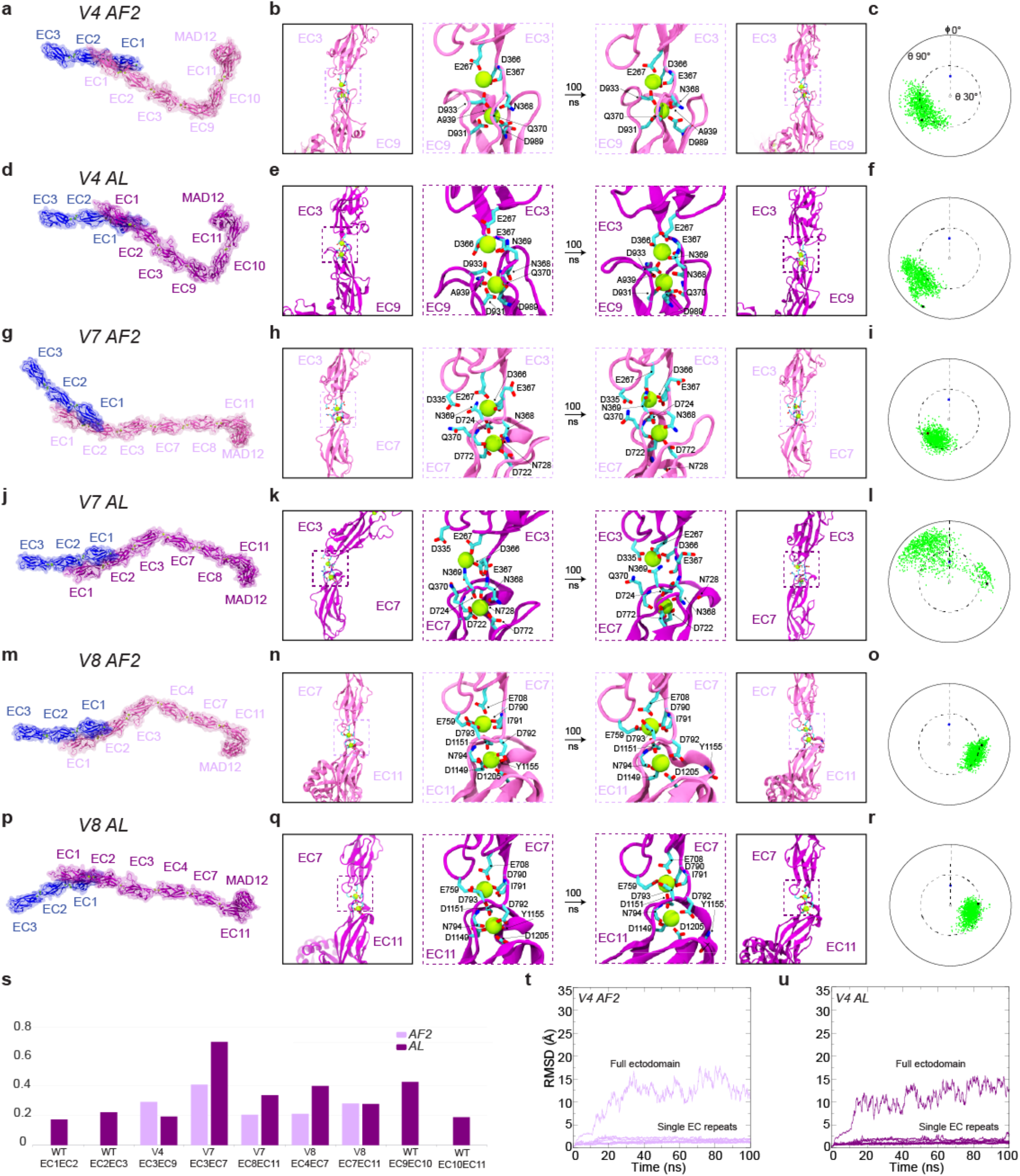
| Structural models and equilibrium MD simulations of mini-PCDH15 proteins. a, Full ectodomain of mini- PCDH15-V4 AF2. **b**, Views of the EC3-EC9 linker region at beginning and end of 100-ns MD equilibrations. **c**, Inter-repeat linker flexibility of EC3-EC9 repeats in mini-PCDH15-V4 AF2. Flexibility was quantified by aligning the longest principal axis of the EC3 repeat to the *z* axis, computing the principal axes of the EC9 repeat, and plotting the third principal axis projection (green) on the *x*-*y* plane in a polar plot. Tilt (𝜃) and azimuthal angle (𝜙) are indicated. The initial orientations for EC3-EC9 and CDH23 EC1-2 (2WHV; 𝜑 ≡ 0^∘^) are shown as black and blue circles, respectively. **d**, Full ectodomain of mini-PCDH15- V4 AL. **e**, Views of EC3-EC9 in mini-PCDH15-V4 AL (as in **b**). **f**, Inter-repeat linker flexibility (as in **c**) of EC3-EC9 repeats in mini-PCDH15-V4 AL. **g**, Full ectodomain of mini-PCDH15-V7 AF2. **h**, Views of EC3-EC7 in mini-PCDH15-V7 AF2 (as in **b**). **i**, Inter-repeat linker flexibility (as in **c**) of EC3-EC7 in mini-PCDH15-V7 AF2. **j**, Full ectodomain of the mini-PCDH15-V7 AL. **k**, Views of EC3-EC7 in mini-PCDH15-V7 AL (as in **b**). **l**, Inter-repeat linker flexibility (as in **c**) of EC3-EC7 in mini- PCDH15-V7 AL. **m**, Full ectodomain of mini-PCDH15-V8 AF2. **n**, Views of EC7-EC11 in mini-PCDH15-V8 AF2 (as in **b**). **o**, Inter-repeat linker flexibility (as in **c**) of EC4-EC7 in mini-PCDH15-V8 AF2. **p**, Full ectodomain of mini-PCDH15-V8 AL. **q**, Views of EC7-EC11 in mini-PCDH15-V8 AL (as in **b**). **r**, Inter-repeat linker flexibility (as in **c**) of EC7-EC11 in mini-PCDH15- V8 AL. **s**, Standard deviation of distances between linker region terminal residues computed during equilibrations. **t-u**, RMSD as a function of time for the full ectodomains and individual EC repeats of the mini-PCDH15-V4 AF2 and AL models. All molecular images display protein backbones in cartoon and sidechains in sticks. Mini-PCDH15 proteins with AF2 predicted engineered linker regions are shown in mauve, whereas those with linker regions assembled based on alignments (AL) are in purple.

Equilibrium MD simulations for the six models allowed us to evaluate stability (RMSD), flexibility (relative orientation between consecutive EC repeats), and linear rigidity (standard deviation of distances between linker region terminal residues) of engineered EC-EC linkers **(Fig. 4, S1,** and **S2;** *see Supplementary Materials for details*). Overall, stability of individual EC repeats was maintained during the simulations **(Fig. 4t** and **4u, S2)**. Simulation analyses revealed low flexibility for the engineered EC7-EC11, moderate flexibility for the engineered EC3-EC9, EC7-EC11, and EC8-EC11 linkers, and a mix of moderate and high flexibility for EC3-EC7 for AF2 and AL models **(Fig. 4a-r)**. Linear rigidity values of engineered EC-EC linkers were compared to both native canonical (EC1-2 and EC10-11) and native non-canonical linkers (EC2-3 and EC9-10). Intriguingly, the EC3- EC7 connection of mini-PCDH15-V7 emerged as the least rigid **(Fig. 4s)**, suggesting that this would be the most extensible linker.

The equilibrated models for mini-PCDH15-V4, -V7, and -V8 served as starting configurations for SMD simulations probing their elasticity. We conducted constant-velocity SMD simulations of their complexes with CDH23 EC1-3 at three varying speeds^46,47^: 10 nm/ns, 1 nm/ns, and 0.1 nm/ns for each model **(Fig. 5** and **S3)**. Results for the mini-PCDH15-V4 + CDH23 EC1-3 complexes, particularly at 0.1 nm/ns, suggest a mechanical response that closely resembles the WT PCDH15 EC1-MAD12 + CDH23 EC1-3 *in silico* (Movies 2 and 3). Both the AF2 and AL models of mini-PCDH15-V4 displayed a first soft phase akin to WT PCDH15, characterized by unbending of the Ca^2+^-free EC9-10 linker region and comparable soft effective spring constants (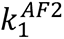 ∼ 3.3 mN/m and ∼ 9 nm extensibility for the mini-PCDH15-V4 AF2 model; 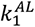∼ 3.3 mN/m and ∼ 7 nm extensibility for the mini-PCDH15-V4 AL model, 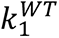∼ 3.3 mN/m and ∼ 10 nm extensibility) as depicted in **(Fig. 5b; Table S3, S3; Sim1d; Sim2d)**. The soft phase was followed by an extension of the EC2-3 and EC9-10 linker regions and then the stretching of the entire protein chain (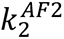∼ 34.2 mN/m and ∼ 5 nm extensibility for the mini-PCDH15-V4 AF2 model; 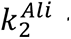 ∼ 17.2 mN/m and ∼ 7.5 nm extensibility for the mini-PCDH15-V4 AL model; 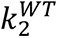∼ 24.4 mN/m and ∼ 5 nm extensibility for the WT PCDH15 ectodomain; **Fig. 5b; Table S2; Sim1d; Sim2d)**. The subsequent unrolling and unfolding of the MAD12 domain occurred at force peaks of ∼ 276 pN for the mini- PCDH15-V4 AF2 model and ∼ 253 pN for the mini-PCDH15-V4 AL model **(Fig. 5g** and **Table S2, Movies 2 and 3).** During these stretching simulations we did not observe unbinding of CDH23 EC1-3 from the mini-PCDH15- V4 protein. Simulations of mini-PCDH15-V7 and mini-PCDH15-V8 bound to CDH23 EC1-3 stretched at 0.1 nm/ns showed that these variants are more rigid than mini-PCDH15-V4. This increased rigidity is attributed to the absence of a soft elasticity phase **(Movies 4 to 7)** and the presence of rigid engineered linkers **(Fig. S4).** Specifically, the mini- PCDH15-V7 AF2 and AL models combined with CDH23 EC1-3 had effective spring constants of ∼ 36.3 mN/m and ∼ 33.0 mN/m with extensibilities of ∼ 4.5 nm and ∼ 5.5 nm, respectively (**Fig. 5d**, *upper* panel; **Table S3; Sim3d)**. The mini PCDH15-V8 AF2 and AL models combined with CDH23 EC1-3 had even higher effective spring constants (∼ 53.9 mN/m and ∼ 55.5 mN/m) than those of the second stretching phase of WT PCDH15 with extensibilities of ∼ 3.5 nm and ∼ 4.0 nm, respectively **(Fig. 5e,f; Table S2; Sim5d** and **Sim 6d; Table S3).**

**Fig. 5.**
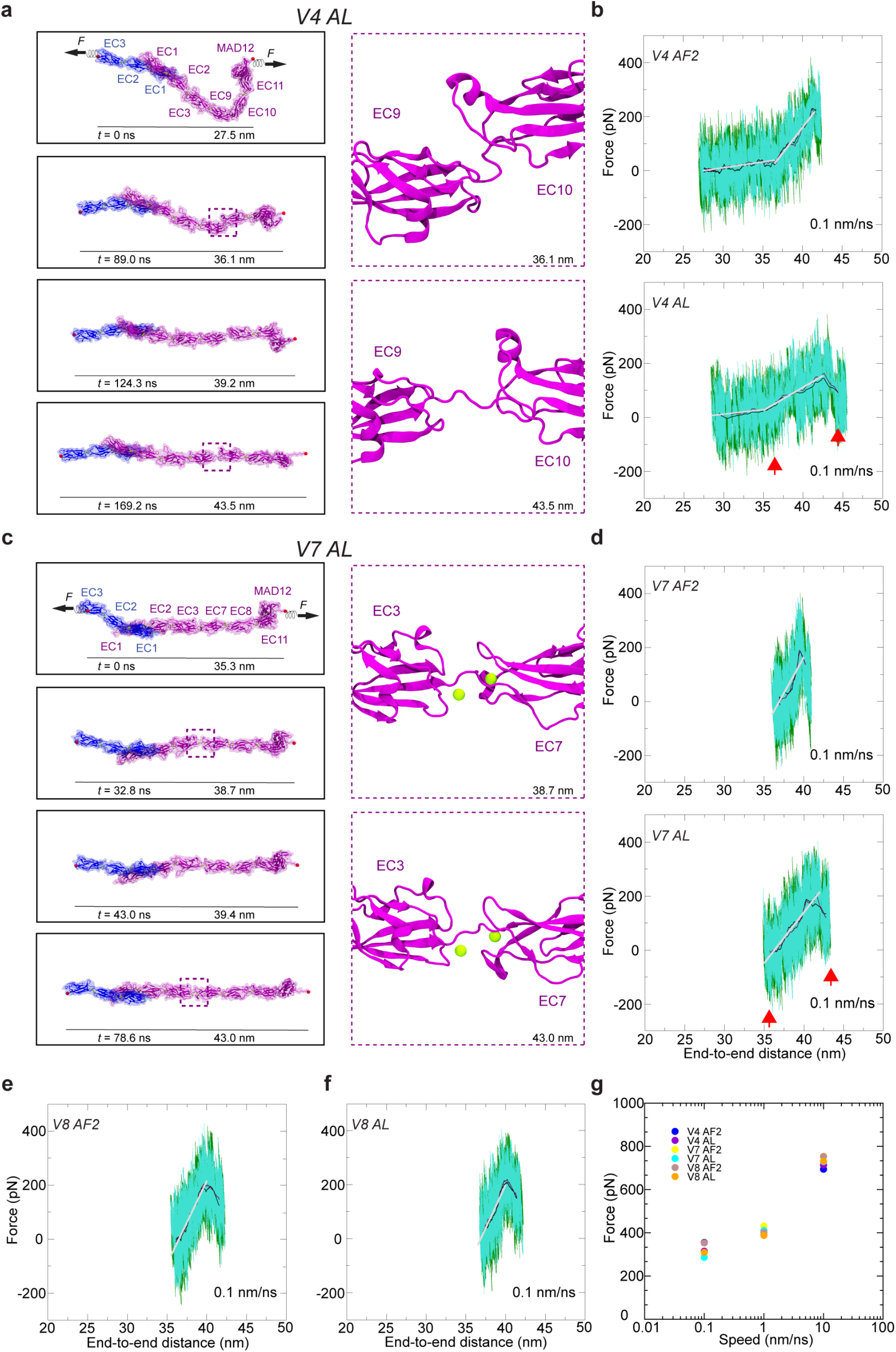
**| Elasticity of monomeric mini-PCDH15 proteins in complex with CDH23 EC1-3. a**, Snapshots of mini-PCDH15- V4 AL + CDH23 EC1-3 during stretching at 0.1 nm/ns (Sim2d). C-terminal C*α* atoms are shown as red spheres. Applied forces are indicated with spring arrows. Insets highlight the most extended EC linker (EC9-EC10). **b**, Force versus end-to-end distance for constant-velocity stretching of mini-PCDH15-V4 AF2 (top) and AL (bottom) + CDH23 EC1-3 at 0.1 nm/ns. Forces at both ends are in dark green and turquoise, with 10-ns running averages in black and maroon. Gray lines are fits for elasticity analysis. **c**, Snapshots mini-PCDH15-V7 AL + CDH23 EC1-3 during stretching at 0.1 nm/ns (Sim4d). **d**, Force versus end-to-end distance for constant-velocity stretching of mini-PCDH15-V7 AF2 (top) and AL (bottom) + CDH23 EC1-3. **e** & **f**, Force versus end-to-end distance for mini-PCDH15-V8 AF2 (**e**) and AL (**f**) models**. g**, *In silico* force peak maxima versus stretching speed for mini-PCDH15 proteins in complex with CDH23 EC1-3.

Collectively, our results suggest that mini-PCDH15-V8 is the most rigid version with minimal extensibility, which correlates with its inability to restore hearing in our previous *in vivo* gene therapy experiments^51^. Conversely, mini-PCDH15-V4 was predicted to be the most flexible and to have the most WT-like mechanical response, providing a possible explanation for why it was the optimal version that restored hearing in *Pcdh*15-KO mice in our previous *in vivo* gene therapy experiments.

### NanoDSF shows differential thermal stabilities for mini-PCDH15 ectodomains in solution

Next, we investigated the impact of EC deletions and engineered EC-EC linker regions on mini-PCDH15’s protein stability. We carried out label-free nanoscale differential scanning fluorimetry (NanoDSF) experiments using the mini-PCDH15-V4, -V7, and -V8 ectodomains as well as the WT PCDH15 EC1-MAD12 ectodomain as a reference **(Fig. 6)**. NanoDSF thermal folding analysis showed that mini-PCDH15s are stable above body temperature (> 37 °C), with thermal unfolding beginning at (*T*onset) 46.0 °C, 42.5 °C and 44.0 °C, respectively **(Fig. 6)**. Interestingly, the WT PCDH15 EC1-MAD12 ectodomain was more stable with *T*onset ∼ 55.1 °C. In addition, we determined the thermal unfolding transition midpoints (*T*m — melting temperature or point at which 50% of the protein is unfolded) for all three mini-PCDH15-V4, -V7, -V8, and WT ectodomains, which were ∼ 65.4 °C, ∼ 62.1 °C, ∼ 62.0 °C, and 72.5 °C, respectively **(Fig. 6)**. These results indicate that all mini-PCDH15 ectodomains are less stable than the WT, with differences in *T*m values that correlate with hearing rescuing ability. Subtle differences in thermal stability might be another reason why mini-PCDH15-V4, but not -V7 and -V8, rescue hearing.

**Fig. 6.**
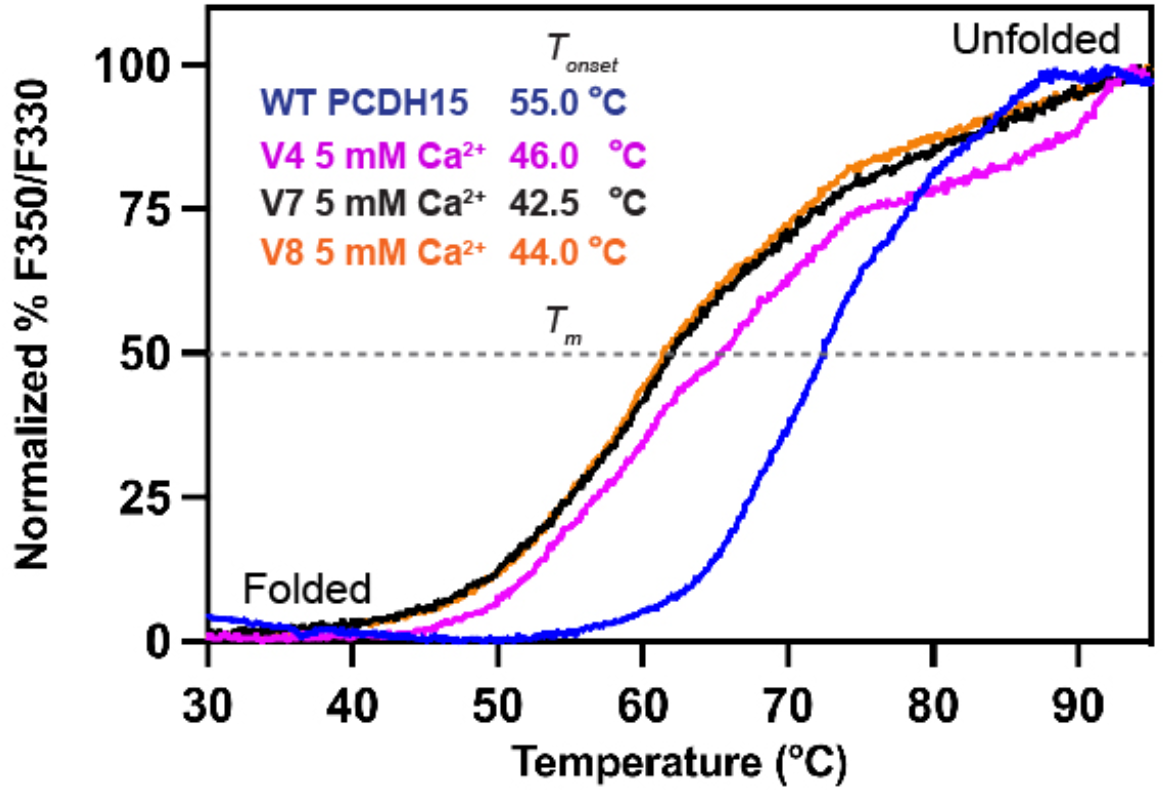
**| Thermal stability of engineered mini-PCDH15s**. Normalized % F350/F330 unfolding curves for mini-PCDH15-V4 (magenta), -V7 (maroon), and -V8 (orange) ectodomains compared to WT PCDH15 EC1-MAD12 (blue). Samples show thermal transitions from folded to unfolded above 42 °C (*T*onset). Dashed line represents the melting temperature point (*Tm*).

## DISCUSSION

Very few mini-genes have been used for AAV-based gene therapy and only a handful of clinical trials are ongoing worldwide^56^. To the best of our knowledge, the first AAV-mini-gene strategy was reported for the cystic fibrosis transmembrane conductance regulator (CFTR) protein in 1998^57^, showing that mini-CFTR membrane proteins could mimic the open probability, ion permeation, and time- and voltage-independence properties of the native channel^58–60^. Other examples of AAV mini-genes are the natural mini-dysferlins^61^, human mini-dystrophins^62^ and their micro-dystrophin variants^41,62–69^, and mini-otoferlins^70^. Although most of these minigenes have not been used in the clinic, the US federal drug administration (FDA) recently approved the AAV-micro-dystrophin treatment, SRP-9001^71^. These recent developments further inspire structure-guided protein shortening approaches offering development of promising functional mini-gene proteins.

In the case of PCDH15, we used structural and biophysical information about its ectodomain to design mini- PCDH15 variants with 3 to 5 EC repeats removed^46,51^. We preserved domains involved in parallel dimerization (EC2-3 and EC11-MAD12) and interactions with CDH23 (EC1-2) and preserved native EC repeat sequences by creating engineered non-native EC-EC linker regions. Three mini-PCDH15 variants (V4, V7, and V8) showed distinct efficacy in rescuing hearing in mutant mice^51^: mini-PCDH15-V4 rescued hearing well, while mini-PCDH15-V7 partially rescued hearing and mini-PCDH15-V8 did not. Here, we have used various techniques to find the factors that determine this differential hearing rescue efficacy. We show that all three mini-PCDH15s can mediate hair-cell mechanotransduction *in vitro,* thus ruling out issues with protein expression or trafficking. SEC- MALS data^51^, negative stain 2D-class averages, and X-ray crystallography data corroborate that our engineered EC-EC linker regions are properly folded and preserve mini-PCDH15 dimerization. We had also shown that all three mini-PCDH15s were able to interact with CDH23^51^. All these results imply that more subtle differences among mini-PCDH15s are the cause of the differential efficacy in hearing rescue.

Interestingly, SMD simulations indicate that the elasticity of monomeric mini-PCDH15-V4 closely resembles that of WT PCDH15, primarily due to the presence of a bent EC9-10 linker. Conversely, monomeric mini-PCDH15- V7 and -V8 are predicted to be more rigid. When we evaluated the thermal stability of mini-PCDH15s using NanoDSF with saturating amounts of Ca^2+^, we found that mini-PCDH15-V4 is slightly more stable than mini- PCDH15-V7 and mini-PCDH15-V8. These differences may be more drastic at the lower, physiological Ca^2+^ concentrations of the inner ear^72^. Based on these results, we suggest that elasticity and thermal stability are key determinants of mini-PCDH15 efficacy in rescuing hearing. Since little is known about PCDH15’s function in the retina, it is hard to speculate whether these differences will affect their ability to rescue the vision loss.

Our results and predictions indicate that any new mini-PCDH15 must have the EC9-10 repeats to provide the right elasticity, in addition to points of dimerization (PCDH15 EC2-3 and EC11-MAD12) and the domain involved in interactions with CDH23 (PCDH15 EC1-2). These constraints do not leave many options for alternates to mini- PCDH15-V4 that can fit in AAVs, except perhaps for an even shorter mini-PCDH15 with an ectodomain made of EC1-EC2-EC3-EC10-EC11-MAD12, where the EC3-EC10 linker region could be engineered to behave as EC9-10. In parallel, improvements for mini-PCDH15-V4 might target thermal stability – the WT ectodomain is significantly more stable than any of the mini-PCDH15s, so utilizing strategies that can increase melting temperature could further improve the performance of mini-PCDH15s in rescuing sensory function. These strategies may include increasing affinity for Ca^2+^, a known stabilizer of cadherin ectodomains that is found at low concentrations in the inner ear.

The apparent necessity of the flexible EC9-10 repeats to rescue hearing in mice using mini-PCDH15s also provides hints about the role played by tip links in inner-ear mechanotransduction. Since the development of the gating spring theory for hair-cell mechanotransduction^73,74^, where an elastic element with a spring constant of ∼ 1 mN/m pulls on the channel, tip links have been put forward as potential gating springs^75^. Yet CDH23 EC repeats are stiff^76^, and the role played by PCDH15 largely depends on the resting tension, as this multimodal protein with a non-linear elastic response can behave as both a soft and stiff spring depending on Ca^2+^ concentration, oligomerization state, and level of stretching^46^. The molecular identity of the gating spring remains unclear, but our results indicate that the elasticity of PCDH15 is important for sustained hearing and tip-link function.

Our results also have implications beyond inner-ear mechanotransduction. PCDH15 is found in retinal photoreceptors, where mutations in it cause progressive degeneration leading to blindness. Since all mini-PCDH15s tested here can interact with CDH23 and rescue hair-cell transduction, form dimers, and fold properly, it is possible that all three will work well in the presumably less mechanically demanding environment of photoreceptors. The design approach used to create and test mini-PCDH15s^51^ could also be applied to other members of the cadherin superfamily of proteins involved in disease.

## METHODS

### Cloning, expression, and purification of bacterially expressed engineered EC-EC fragments

Mouse PCDH15 EC3-EC7 and EC4-EC7 were subcloned into *NdeI/NheI* and *XhoI* sites of the pET21a+ vector. These constructs were used for protein expression in Rosetta (DE3) competent cells (Novagen) cultured in TB as reported previously^11,46,47^. Cell cultures were grown at 37 °C and induction was started at OD600 = 0.4 - 0.6 with 1 mM of IPTG. Induction was done at 30 °C for ∼16 hr. Cells were pelleted at 4 °C and resuspended in denaturing buffer (20 mM TrisHCl [pH 7.5], 6 M guanidine hydrochloride, 10 mM CaCl2, and 20 mM imidazole). The cell mix was lysed by sonication. The cleared lysates were pelleted at 18,000 rpm over 40 min and the supernatant was incubated with Ni-Sepharose beads (GE Healthcare) for 1 h. The mix was centrifuged for 5 min to pellet beads. Supernatants were collected and the beads were washed with denaturing buffer. The process was repeated 3 times. Finally, the protein was eluted with denaturing buffer supplemented with 500 mM imidazole. The PCDH15 EC3-EC7 protein fragment was refolded overnight at 4 °C by dialysis of 50 mL of eluted denatured protein at 0.5 mg/mL into 1,000 mL of refolding buffer (20 mM TrisHCl [pH 8.0], 150 mM KCl, 5 mM CaCl2, 400 mM L-arginine, 1 mM of glutathione oxidized). Similarly, PCDH15 EC4-EC7 was refolded overnight using 20 mM TrisHCl [pH 8.0], 150 mM KCl, 5 mM CaCl2, and 400 mM L-arginine. Samples were vigorously stirred overnight. Both proteins were concentrated to a 2 mL volume using Amicon Ultra-15 filters at 3,500 rpm with mixing every 20 min. Refolded proteins were further purified on a Superdex-200 16/600 column (GE Healthcare) using 20 mM TrisHCl [pH 8.0], 150 mM KCl, and 5 mM CaCl2 using an ÄKTA Purifier system at 4 °C. Purity of the constructs were analyzed by SDS-PAGE and pure fractions were concentrated by centrifugation using Amicon Ultra-15 to 6-8 mg/mL for crystallization trials at 4 °C. Protein concentrations were determined using the NanoDrop (Thermo Scientific).

### Cloning, expression, and purification of mammalian-expressed mini-PCDH15s

The coding sequence of *mm* mini-PCDH15 ectodomains were subcloned into *NdeI* and *XhoI* sites of a CMV vector. The native signal sequence was preserved at the N-terminal end and a double-hexahistidine tag was inserted at the C-terminal end of the protein sequence. Proteins were expressed in suspension Expi293F cells at 37 °C and purified under native conditions with TALON metal affinity resin beads (Takara) and SEC experiments, as reported^51^.

### Nanoscale differential scanning fluorimetry (NanoDSF)

Purified proteins after SEC were concentrated to 0.5-1 mg/mL for NanoDSF using a Prometheus NT.48 (Nanotemper). Capillaries were filled up with ∼ 15 µL of protein for melting scans between 20-95 °C with a pre-stabilization phase of 1 min and a temperature slope of 2 °C/min (37 min, 37 s total scanning time). Data were processed using a PR.ThermControl v2.1.2 software and plotted using the Prism software. NanoDSF experiments were done with at least one duplicate and temperatures were averaged. For protein folding analysis, the normalized % F350/F330 ratio of fluorescent intensities were plotted. The temperature at which the protein begins to unfold (*T*onset) was determined with the first inflection point at which the % F350/F330 intensities increased. Thermal unfolding transition midpoints (*T*m) were determined as the temperature values at 50% F350/F330 ratio intensities.

### Protein crystallization

Proteins were purified using a SEC buffer containing 20 mM TrisHCl [pH 8.0], 150 mM KCl, and 5 mM CaCl2. Initial protein crystal hits for PCDH15 EC3-EC7 were obtained using an initial volume of 75 μL of reservoir solution (0.1 M Tris.HCl pH 8.5 and 15% ^W^/V PEG 20,000) in a sitting-drop vapor diffusion setup at 4 °C. The crystallization droplets were composed of equal volumes (0.6 μL) of protein solution (at 8.0 mg/mL) and reservoir solution. Final high-resolution diffracting crystal were refined by adding a maximum concentration of 0.6 μM of spermine tetrahydrochloride in the reservoir solution and mixed with equal amount of protein (at 8.0 mg/mL) and reservoir solution (0.6 μL) as indicated above. For the PCDH15 EC4-EC7 protein, crystals were obtained in 0.1 M sodium cacodylate pH 6.5, 40% ^V^/V MPD, 5% ^W^/V PEG 8000 as a reservoir solution. Protein (0.5 μL at 6.6 mg/mL) and reservoir solution (0.6 μL) were mixed for crystallization.

### Data collection and structure determination

All crystals were cryoprotected in reservoir solution plus 25% ^V^/V MPD and were cryo-cooled in liquid N2. X-ray diffraction data is reported in Table 1 and was processed with HKL2000^77^. The final structures were determined by molecular replacement using PHASER^78^. For PCDH15 EC3-EC7, an initial search was based on the homology model of *Homo sapiens* PCDH15 EC3 based on (PDB ID: 54TM)^79^ and a separate EC7 structure as a template (PDB ID: 5TPK)^46^. For PCDH15 EC4-EC7, the initial search with PHASER^78^ was carried out using individual EC4 and EC7 repeats as templates (PDB ID: 5W1D). All molecules were refined using COOT^80^ and REFMAC5^81^ and restrained TLS refinement was used for PCDH15 EC4-EC7. Data collection and refinement statistics are provided in **Table 1**.

### Structural models for simulations

We assembled models of the full ectodomain of monomeric mini-PCDH15 proteins by integrating crystal structures of PCDH15^46^ along with engineered linker regions. The crystal-structure-based model of full-length PCDH15 EC1-MAD12 bound to CDH23 EC1-3 presented by Choudhary et al^46^ was used as a starting base model to build all mini-PCDH15 ectodomains. Two strategies were used to obtain the engineered linker regions. In the first one, AF2^82^ was used to predict them. These predictions were done without using a structural template in a Colab notebook (ColabFold v1.5.5, Google research)^83^. Specifically, for mini-PCDH15-V4 AF2, we predicted the structure of engineered EC3-EC9 in AF2 by using a combination of sequences: p.D236 to p.Q370 (covering

PCDH15 EC3 and linkers) followed by p.P902 to p.E1010 (covering PCDH15 EC9 and linkers). We then aligned C*α*atoms of the C-terminal end of EC2 in our base model (p.236DGDDL) to C*α* atoms of the N-terminal end of our AF2-predicted EC3-EC9 domain, deleted the AF2-predicted p.236DGDDL, and kept the base model up to EC2 with the EC3-EC9 linker region integrated **(Fig. S5)**. Similarly, we aligned C*α* atoms of the N-terminal end of PCDH15 EC10 in our base model to the C-terminal end of the AF2-predicted EC3-EC9 domain (p.1006LHPGE), deleted the AF2-predicted p.1006LHPGE, and kept the base model from EC10 on. Finally, the aligned regions of mini-PCDH15-V4 AF2 **(Fig. S5)** were regularized in COOT. A similar process was consistently used across all AF2-based versions of mini-PCDH15 proteins (mini-PCDH15-V7 AF2 with AF2 predictions for p.D236 to p.Q370 covering EC3 and linkers, p.A693 to p.Y901 covering EC7-9 and linkers, and p.P1120 to p.1226 covering EC11 and linkers; mini-PCDH15-V8 AF2 with AF2 predictions for p.D366 to p.N484 covering EC4 and linker, p.A693 to p.N792 covering EC7 and linkers, and p.1120 to p.1226 covering EC11 and linkers).

The second approach used to build mini-PCDH15 ectodomains involved aligning parts of our base model (full-length PCDH15 EC1-MAD12 bound to CDH23 EC1-3) and building the engineered linkers manually by connecting the N- and C-termini of EC repeats in COOT. For the mini-PCDH15-V4 ectodomain with the engineered linker EC3-EC9, we utilized the base model up to PCDH15 EC3 and including the EC3-4 connecting linker sequence (p.366DENNQ; **Fig. S6).** This assembly was then connected to the base model starting from PCDH15 EC9 **(Fig. S6,** in lime) by aligning the C*α* atoms of the last two residues (p.369NQ) from the EC3-4 linker p.366DENNQ to the C*α* atoms of the last two residues (p.900DY) from the EC8-9 linker p.897DMNDY. Finally, we deleted p.897DMNDY and regularized the engineered linker region in COOT. A similar process was consistently used across all alignment-based versions of mini-PCDH15 proteins (mini-PCDH15-V7 AL and mini- PCDH15-V8 AL). The six-resulting mini-PCDH15 models were labeled mini-PCDH15-V4 AF2, mini-PCDH15-V4 AL, mini-PCDH15-V7 AF2, mini-PCDH15-V7 AL, mini-PCDH15-V8 AF2, and mini-PCDH15-V8 AL. Engineered linker regions in all versions were modeled with two Ca^2+^ ions at sites 2 and 3. All alignments used to build AF2 and AL models were done using COOT.

We also utilized AF3 to predict the structures of our engineered linkers. The RMSD value differences between the AF2 and AF3 predicted structures of our engineered linkers were minimal **(Table S4)**, validating the use of AF2 and AF3 as equally accurate prediction tools.

### Molecular dynamics simulations

Hydrogen atoms were appended to all protein structures and models using the psfgen tool. Simulation systems were prepared with VMD plugins—autopsfgen for structure setup, solvate for system hydration, orient for optimal positioning, and autoionize for ion placement—resulting in proteins solvated in explicit water with randomly distributed ions to emulate a physiological endolymph environment characterized by 150 mM KCl. MD simulations were conducted using versions 2.12 and 2.13 of NAMD^84^, employing the CHARMM36 force field with the CMAP correction for proteins and the TIP3P water model^85^. Van der Waals interactions were computed with a 12 Å cutoff, incorporating a switching function beginning at 10 Å. We used periodic boundary conditions and the particle mesh Ewald method for the calculation of long-range electrostatic forces, ensuring a grid point density exceeding 1 Å^-3^. The simulations adopted a uniform integration time step of 2 fs with the SHAKE algorithm maintaining bond length constraints. Constant pressure simulations (*NpT*) at 1 atm were conducted using the hybrid Nosé-Hoover Langevin piston method with a 200 fs decay period and a 100 fs damping time constant. Temperature control at *T* = 310 K was achieved through Langevin dynamics, ensuring constant temperature conditions throughout the simulations. Each system was equilibrated in various steps. A 1,000-step minimization was followed by a 0.1-ns long simulation with backbone restrained and a 1-ns long step of free dynamics with a Langevin damping coefficient of 1 ps^-1^. Lastly, we performed 100 ns of free dynamics with a Langevin damping coefficient of 0.1 ps^-1^.

Each simulation system underwent energy minimization and was equilibrated under the *NpT* ensemble, preparing it for further equilibrium and SMD simulations. For constant-velocity stretching simulations, we employed the SMD technique in conjunction with the NAMD Tcl forces interface^86–89^. This involved anchoring the Cα atoms of selected terminal residues to virtual springs, each with a stiffness of 1 kcal mol^-1^ Å^-2^, allowing for controlled manipulation. The opposite ends of these springs were then extended away from the protein at a constant velocity, simulating mechanical stretching. The orientation for stretching was aligned along the *x*-axis, corresponding to the vector that connects the terminal regions of the proteins within the simulated complexes.

### Simulation analysis

Applied forces during the simulations were computed by using the extension observed in the virtual springs attached to the protein. We assessed the protein’s mechanical properties by performing linear regression analyses on the force versus end-to-end distance data, facilitating the calculation of protein stiffness. To mitigate the impact of local fluctuations, maximum force peaks along with their averages were derived from 50-ps running averages. In the constant-velocity SMD simulations for systems Sim1d, Sim2d, Sim3d, Sim4d, Sim5d, and Sim6d, the end-to-end distance was quantified by measuring the separation between the centers-of-mass of Cα atoms at opposing ends of the protein. The principal axes of EC repeats were identified using the Orient plugin in VMD, assisting in the analysis of protein orientation and dynamics. Graphical representations and linear curve fitting were done using Xmgrace. VMD^90^ was utilized to generate molecular visualizations.

### Viral vector production

AAV cDNA plasmids encoding mini-PCDH15-V7 and mini-PCDH15-V8 were obtained from Addgene, accession numbers 199186 and 199187, respectively. AAVs for this study were produced by PackGene, Houston, TX. Serotype AAV9-PHP.B vectors were packaged in HEK293T cells using polyethylenimine-mediated co-transfection of the pAAV transfer plasmid, pHelper plasmid, and RepCap plasmid pUCmini-iCAP-PHP.B. The media and cells were harvested approximately 120 hours post-transfection. AAV9-PHP.B viruses were released and subjected to discontinuous density iodixanol (OptiPrep, Axis-Shield) gradient ultracentrifugation. The AAV vector-containing iodixanol fractions were isolated, concentrated by diafiltration, and the purified AAV vectors were quantified by Q-PCR. The final titer for AAV encoding mini-PCDH15-V7 was 2.09E+13 GC/mL and for mini- PCDH15-V8 was 2.00E+13 GC/mL. Vectors were aliquoted for single-use and stored at −80°C until needed.

### Round window membrane injection of AAV-mini-PCDH15 in neonatal mice

All animal procedures were approved by the Mass Eye and Ear Institutional Animal Care and Use Committee. The round window membrane (RWM) injections were performed under a stereomicroscope. P0-1 *Pcdh15^-/-^* mice and their heterozygote littermates were anesthetized using cryoanesthesia and maintained on an ice pack during the entire procedure. Injections through the RWM were conducted as previously described^51^. A small incision was made under the external ear, enlarged to push aside soft tissues and expose the bulla. The round window niche was visually localized, and 1 µL of viral vector solution was injected at a rate of 60 nL/min using a Nanoliter 2000 Injector (World Precision Instruments). The surgical incision was sutured using 7-0 Vycril thread, with standard postoperative care provided following the procedure.

### Cochlear explant FM1-43 dye loading and fluorescence imaging

On postnatal day 3 (P3), mice of either sex were cryo-anesthetized and euthanized by decapitation. The inner ears were harvested, and the organ of Corti epithelia were acutely dissected in Leibovitz’s L-15 (Invitrogen) cell culture medium. Following dissection, the explants were secured to a coverslip with tungsten pins, then placed in glass-bottom cell culture dish. After gently aspirating the L15 media the explants were incubated for 60 s with FM1-43 (5 µM) in L15. Next, the solution was aspirated, and the excess FM1-43 dye was neutralized using a quenching solution composed of 0.2 mM SCAS (Biotium) in L-15 and taken immediately for live imaging. Imaging was performed using a Leica SP5 confocal microscope equipped with a 40x, 0.8 NA water-dipping objective lens, with the zoom set to 2x, resulting in an effective pixel size of 189*189 nm. The laser power and smart gain settings were consistently maintained across all experimental conditions, although animals from different litters were imaged on different days.

### Negative stain EM sample preparation and analysis

For each mini-PCDH15 variant, a dilution series of the concentrated stock solution was prepared ranging from 1:10 to 1:1000. A small drop (2.5 mL) of the protein solution was applied to the carbon side of a glow-discharged grid (EMS, CF300-CU) and left on the grid for 1 min. The grid was blotted from the side with a piece of filter paper without allowing the surface to completely dry. Excess protein was rinsed off by touching a 20 mL drop of MQ water and briefly blotting – this was repeated twice. The particles were then stained by touching a 20 mL drop of 2% uranyl acetate, briefly blotting, and repeating these steps with a final 30 second incubation in the uranyl acetate solution. The grid was fully dried and stored for subsequent TEM imaging. Grids were imaged on a FEI Tecnai F20 TEM operated at 200 kV using a Gatan Orius CCD camera and SerialEM for automated image collection. The grid with ideal particle homogeneity and density from each dilution series was used for further analysis.

### Negative stain EM data processing

All negative stain EM images were processed using cryoSPARC (Structura) and followed the same general procedure. CTF estimation was completed using CTFFIND4, after which all images were inspected and curated. Approximately 1,000 particles were manually selected from a subset of the collected images and used to generate initial templates. A template picker was then used to select particles from all images, resulting in over 10,000 particles for classification. 2D classification jobs were run and junk particles were removed.

## AUTHOR CONTRIBUTIONS

P.D. carried out protein expression, purification, and NanoDSF of *mm* mini-PCDH15 versions and WT mm PCDH15 EC1-MAD12, as well as SEC-MALS experiments, data analysis. P.D. and J.B. carried out protein expression, purification, and SEC of mini-PCDH15 versions for negative stain experiments, cloned, purified protein, and obtained crystals of mini-PCDH15 EC3-EC7 fragment. P.D and K.M.P cloned, purified protein, obtained crystals of mini-PCDH15 EC3-EC7. P.D, J.B and K.M.P collected X-ray crystallography datasets, solved, and deposited the X-ray crystal structures in the PDB. P.D, J.B, and H.W. generated AlphaFold models. P.D., H.W. M.S. and A.A.I. analyzed data. H.W and M.S. generated experimentally-based models of mini- PCDH15 proteins for SMD simulations, carried out SMD simulations and SMD analyses. Y.N. performed negative stain experiments and analyzed data. M.S. assisted with model building, SMD simulations, training, experiments, and structure refinement. M.V.I. contributed to the design and cloning of vectors for *in vivo* experiments, as well as conceptualization and manuscript editing. E.H. performed inner ear injections. F.Y. carried out FM1-43 dye loading experiments. P.D, H.W, M.S. and A.A.I. prepared the original manuscript draft. P.D, H.W, M.S. and A.A.I. edited the manuscript and incorporated feedback from all authors. A.A.I. M.S. and D.P.C. conceived the study and secured funding. P.D., A.A.I. and M.S. oversaw the project. All authors contributed to the final version of the manuscript.

## ACKNOWLEDGMENTS

We would like to thank Dr. Eric Mulhall for assistance with the cloning of the mini-PCDH15 variants. Simulations were carried out at the Ohio Supercomputer Center (OSC grants PAS1037 and PAA0217 to M.S.) and San Diego Supercomputer Center (ACCESS grant MCB140226 to M.S.). Part of this work is based upon research conducted at the Northeastern Collaborative Access Team beamlines, which are funded by the National Institute of General Medical Sciences from the National Institutes of Health (P30 GM124165). The Eiger 16M detector on the 24-ID-E beam line is funded by a NIH-ORIP HEI grant (S10OD021527). This research used resources of the Advanced Photon Source, a U.S. Department of Energy (DOE) Office of Science User Facility operated for the DOE Office of Science by Argonne National Laboratory under Contract No. DE-AC02-06CH11357. This study was supported by R01DC017166 (NIDCD) to A.A.I. and MPI R01DC020190 (NIDCD) to A.A.I., D.P.C. and M.S.

## DATA AVAILABILITY

The atomic coordinates of *mm* PCDH15 EC3-EC7 and *mm* PCDH15 EC4-EC7 have been deposited in the Protein Data Bank (PDB) under accession numbers PDB 8TON and 8UMZ, respectively. Other atomic coordinates used in this study for comparisons are available from the PDB and indicated in the text. Other data and materials of this study will be made available upon request. Correspondence and requests for materials should be addressed to A.A.I and M.S.

## ETHICS DECLARATIONS

### Competing interests

Authors declare that they have no competing financial and/or non-financial interests in relation to this work.

## SUPPLEMENTARY MATERIALS

### Assignment and equilibrium dynamics of Ca^2+^ bound to engineered linkers in mini-PCDH15 models

Our AlphaFold2 (AF2) and aligned (AL) models of mini-PCDH15 ectodomains included engineered linker regions where we manually placed bound Ca^2+^ ions. Mini-PCDH15-V4 (Fig. 4a**, 4d)** has one engineered EC3-EC9 linker region with the atypical p.366DENNQ linker motif. The native EC3-EC4 linker region with this motif and with Ca²⁺ observed at sites 2 and 3 ^79^ exhibits reduced Ca²⁺ binding ability compared to what is expected for linker regions with the canonical DXNDN Ca^2+^ binding motif and three bound Ca²⁺. Analysis of our EC3-EC9 AF2 and AL models (Fig. 4 **and S3)**, revealed residues arranged in conformations that were compatible with Ca²⁺ binding at sites 2 and 3, where Ca²⁺ ions were placed (Fig. 4b and 4e, *left middle* panels). Interestingly, Ca^2+^ at site 2 was coordinated by p.N369 in the mini-PCDH15-V4 AL model, but not in the mini-PCDH15-V4 AF2 model. Mini- PCDH15-V4 AF2 and AL models showed consistent Ca^2+^ coordination profiles pre- and post-simulation (Fig. 4b and 4e, *right middle* panels).

Mini-PCDH15-V7 (Fig. 4g and 4j) features two engineered linker regions: EC3-EC7 and EC8-EC11. The EC3- EC7 linker region has the non-canonical p.366DENNQ linker motif. Our AF2 model (Fig. 4h, *left middle*), the AL model (Fig. 4k, *left middle)*, and the crystal structure (Fig. 3a) show a Ca^2+^ bound at site 3, as expected. Notably, our crystal structure of the EC3-EC7 fragment has a K^+^ at site 1, with no cation occupancy at site 2 (Fig. 3a). In contrast, in both the AF2 and the AL models, Ca^2+^ ions were placed at site 2, leading to p.D724 coordinating Ca²⁺ ions at sites 2 and 3, as we previously observed in the native EC3-EC4 linker with residue p.D411 (PDB: 5T4M). In both the AF2 and AL models, the side chains of the linker motif segment p.368NNQ are oriented in a configuration consistent with the EC3-4 crystal structure (Fig. 3a, Fig. 4h and 4k, *left middle*). Equilibrium simulations also revealed that in both models for mini-PCDH15-V7 EC3-EC7, the backbone oxygen atom of p.N728 loses coordination with the Ca^2+^ ion at site 3 (Fig. 4h and 4k, *right middle*), inconsistent with the configuration observed in the EC3-EC7 crystal structure (Fig. 3a). In contrast, Ca^2+^ at site 2 moved close to site 1 during the equilibration of the AL model, to a position that is consistent with that observed for K^+^ in the crystal structure (Fig. 3a).

Our models for the engineered linker region EC8-EC11 of mini-PCDH15-V7 with the p.897DMNDY linker motif, display similar Ca^2+^ coordination at sites 2 and 3 pre- and post-simulation (**Fig. S3b** and **S3e**, middle panels). However, the backbone oxygen atom of p.Y1155 loses its coordination with the Ca²⁺ ion at site 3 after equilibration (**Fig. S3b** and **S3e**, *right middle* panels).

Models for mini-PCDH15-V8 and its engineered EC4-EC7 **(Fig. S3g and S3j)** linker region with the canonical p.480DANDN linker motif displayed Ca^2+^ coordination at sites 2 and 3 (**Fig. S3h** and **S3k**, *left* panels), similar to that observed in our crystal structure (Fig. 3b). Post-simulation, however, both models exhibited a notable shift: the backbone oxygen atom of p.N728 moved away from the Ca^2+^ ion at site 3 (**Fig. S3h** and **S3k**, *right middle* panels), accompanied by a general increase in flexibility **(Fig. S3i** and **S3l)**. Furthermore, our AF2 and AL models of the engineered EC7-EC11 linker region with a canonical p.790DIDDN linker motif had identical Ca^2+^ binding configurations (Fig. 4n and 4q). Equilibrium simulations of mini-PCDH15-V8 AF2 and AL models revealed a consistent pattern: the backbone oxygen atom at p.Y1155 disengaged from the Ca^2+^ ion at site 3 in both models (Fig. 4n and 4q, *right middle* panels), although the overall Ca^2+^ coordination remained unchanged pre- and post-simulation.

## Supplementary Figures and Tables

## Movies

**Movie S1**. Animation of 2D class averages of mini-PCDH15-V4 demonstrating the various conformations present.

**Movie S2**. Forced unbending and unrolling in a simulation of the mini-PCDH15 V4 AF2 (EC1-EC2-EC3-EC9-EC10-EC11- MAD12 in mauve) + CDH23 (EC1-3 in blue) model. Stretching of the complex at 0.1 nm/ns (simulation s1d in SI Appendix, Table S1, 0 - 170.9 ns) results in straightening of the mini-PCDH15 V4 AF2 ectodomain with lengthening of the EC2-3 and EC9-10 linker regions. As the simulation progresses the mini-PCDH15 MAD12 began to unfold from its C-terminal end. The mini-PCDH15 MAD12 eventually unrolled away from EC11 while unfolding continued. The CDH23 EC1-3 did not unbind from mini-PCDH15 V4 AF2.

**Movie S3**. Forced unbending and unrolling in a simulation of the mini-PCDH15 V4 AL (EC1-EC2-EC3-EC9-EC10-EC11- MAD12 in purple) + CDH23 (EC1-3 in blue) model. Stretching of the complex at 0.1 nm/ns (simulation s2d in SI Appendix, Table S1, 0 - 189.2 ns) results in straightening of the mini-PCDH15 V4 AL ectodomain with lengthening of the EC9-10 linker region. As the simulation progressed the mini-PCDH15 MAD12 began to unfold from its C-terminal end. The mini-PCDH15 MAD12 eventually unrolled away from EC11 while unfolding continued. The CDH23 EC1-3 did not unbind from mini- PCDH15 V4 AL.

**Movie S4**. Forced unrolling in a simulation of the mini-PCDH15-V7 AF2 (EC1-EC2-EC3-EC7-EC8-EC11-MAD12 in mauve) + CDH23 (EC1-3 in blue) model. Stretching of the complex at 0.1 nm/ns (simulation s3d in SI Appendix, Table S1, 0 - 84.0 ns) results in straightening of the mini-PCDH15-V7 AF2 ectodomains with minimal lengthening of linker regions. As the simulation progressed the mini-PCDH15 MAD12 began to unfold from its C-terminal end. The mini-PCDH15 MAD12s eventually unrolled away from EC11 while unfolding continued. The CDH23 EC1-3 did not unbind from mini-PCDH15 V7 AF2.

**Movie S5**. Forced unrolling in a simulation of the *mm* (mini-PCDH15-V7 Alignment EC1-EC2-EC3-EC7-EC8-EC11-MAD12 in purple) + (CDH23 EC1-3 in blue) model. Stretching of the complex at 0.1 nm/ns (simulation s4d in SI Appendix, Table S1, 0 - 100.6ns) results in straightening of the mini-PCDH15-V7 Alignment ectodomains with minimal lengthening of linker regions. As the simulation progressed the mini-PCDH15 MAD12s began to unfold from their C-terminal ends. The mini- PCDH15 MAD12s eventually unrolled away from EC11 while unfolding continued. The CDH23 EC1-3 did not unbind from mini-PCDH15 V7 Alignment.

**Movie S6**. Forced unrolling in a simulation of the *mm* (mini-PCDH15-V8 AlphaFold2 EC1-EC2-EC3-EC4-EC7-EC11- MAD12 in mauve) + (CDH23 EC1-3 in blue) model. Stretching of the complex at 0.1 nm/ns (simulation s5d in SI Appendix, Table S1, 0 - 84.5ns) results in straightening of the mini-PCDH15-V8 AlphaFold2 ectodomains with minimal lengthening of linker regions. As the simulation progressed the mini-PCDH15 MAD12s began to unfold from their C-terminal ends. The mini-PCDH15 MAD12s eventually unrolled away from EC11 while unfolding continued. The CDH23 EC1-3 did not unbind from mini-PCDH15 V8 AlphaFold2.

**Movie S7**. Forced unrolling in a simulation of the *mm* (mini-PCDH15-V8 Alignment EC1-EC2-EC3-EC4-EC7-EC11-MAD12 in purple) + (CDH23 EC1-3 in blue) model. Stretching of the complex at 0.1 nm/ns (simulation s6d in SI Appendix, Table S1, 0 - 84.1ns) results in straightening of the mini-PCDH15-V8 Alignment ectodomains with minimal lengthening of linker regions. As the simulation progressed the mini-PCDH15 MAD12s began to unfold from their C-terminal ends. The mini- PCDH15 MAD12s eventually unrolled away from EC11 while unfolding continued. The CDH23 EC1-3 did not unbind from mini-PCDH15 V8 Alignment.

**Fig. S1.**
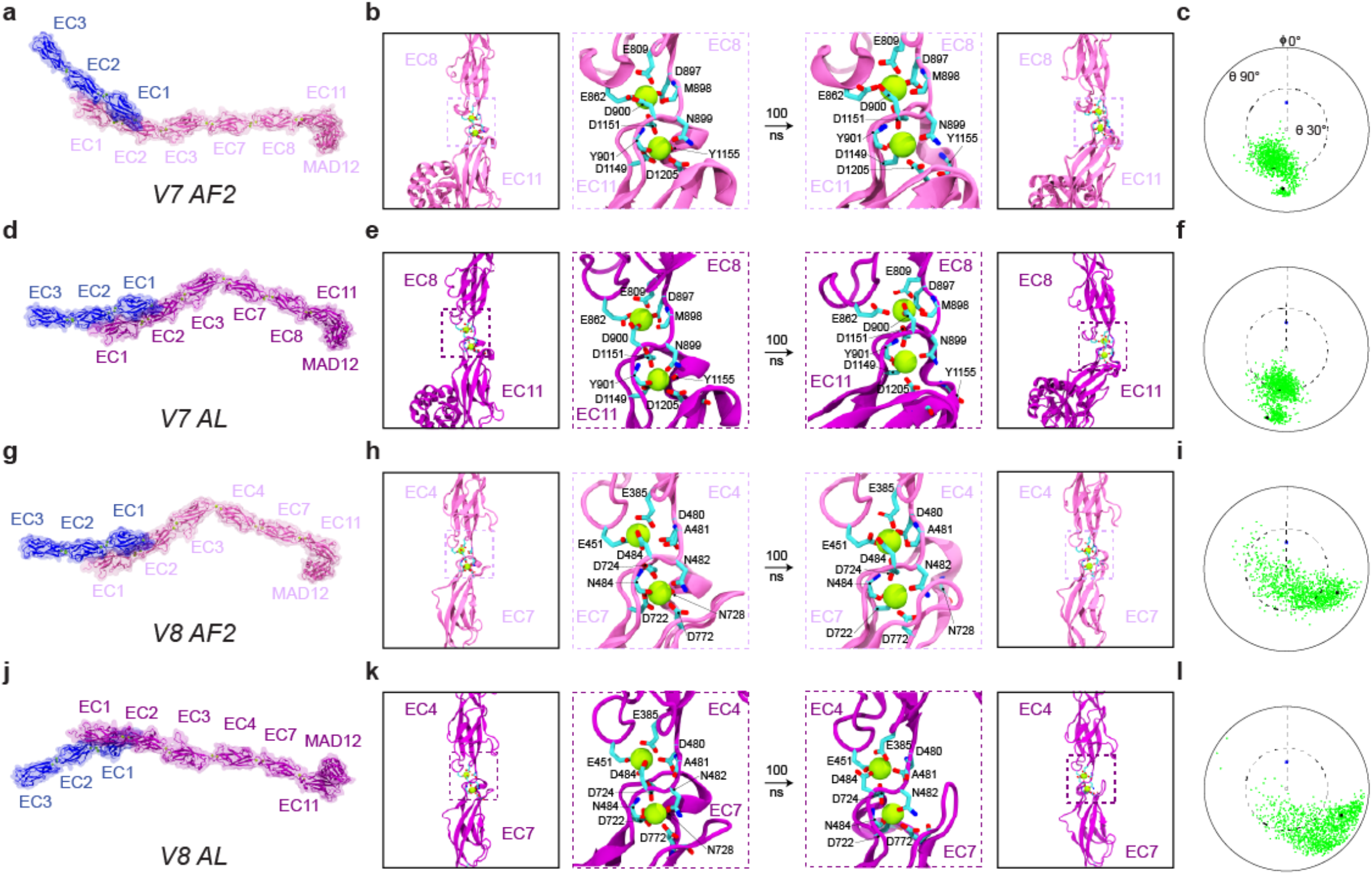
| Structural models and equilibrium MD simulations of mini-PCDH15-V7 and -V8. a, Full ectodomain of mini- PCDH15-V7 AF2. **b**, Views of the EC8-EC11 linker region at beginning and end of 100-ns MD equilibrations. **c**, Inter-repeat linker flexibility of EC8-EC11 repeats in mini-PCDH15-V7 AF2 (as in Fig. 5c). **d,** Full ectodomain of mini-PCDH15-V7 AL. **e**, Views of EC8-EC11 in mini-PCDH15-V7 AL (as in Fig. 5b). **f**, Inter-repeat linker flexibility (as in Fig. 5c) of EC8-EC11 repeats in mini-PCDH15-V7 AL. **g**, Full ectodomain of mini-PCDH15-V8 AF2. **h**, Views of EC4-EC7 in mini-PCDH15-V8 AF2 (as in Fig. 5b). **i**, Inter-repeat linker flexibility (as in Fig. 5c) of EC4-EC7 in mini-PCDH15-V8 AF2. **j**, Full ectodomain of the mini-PCDH15-V8 AL. **k**, Views of EC4-EC7 in mini-PCDH15-V8 AL (as in Fig. 5b). **l**, Inter-repeat linker flexibility (as in Fig. 5c) of EC4-EC7 in mini-PCDH15-V8 AL. All molecular images display protein backbones in cartoon and sidechains in sticks. Mini-PCDH15 proteins with AF2-predicted engineered linker regions are shown in mauve, whereas those assembled based on alignments (AL) are in purple.

**Fig. S2.**
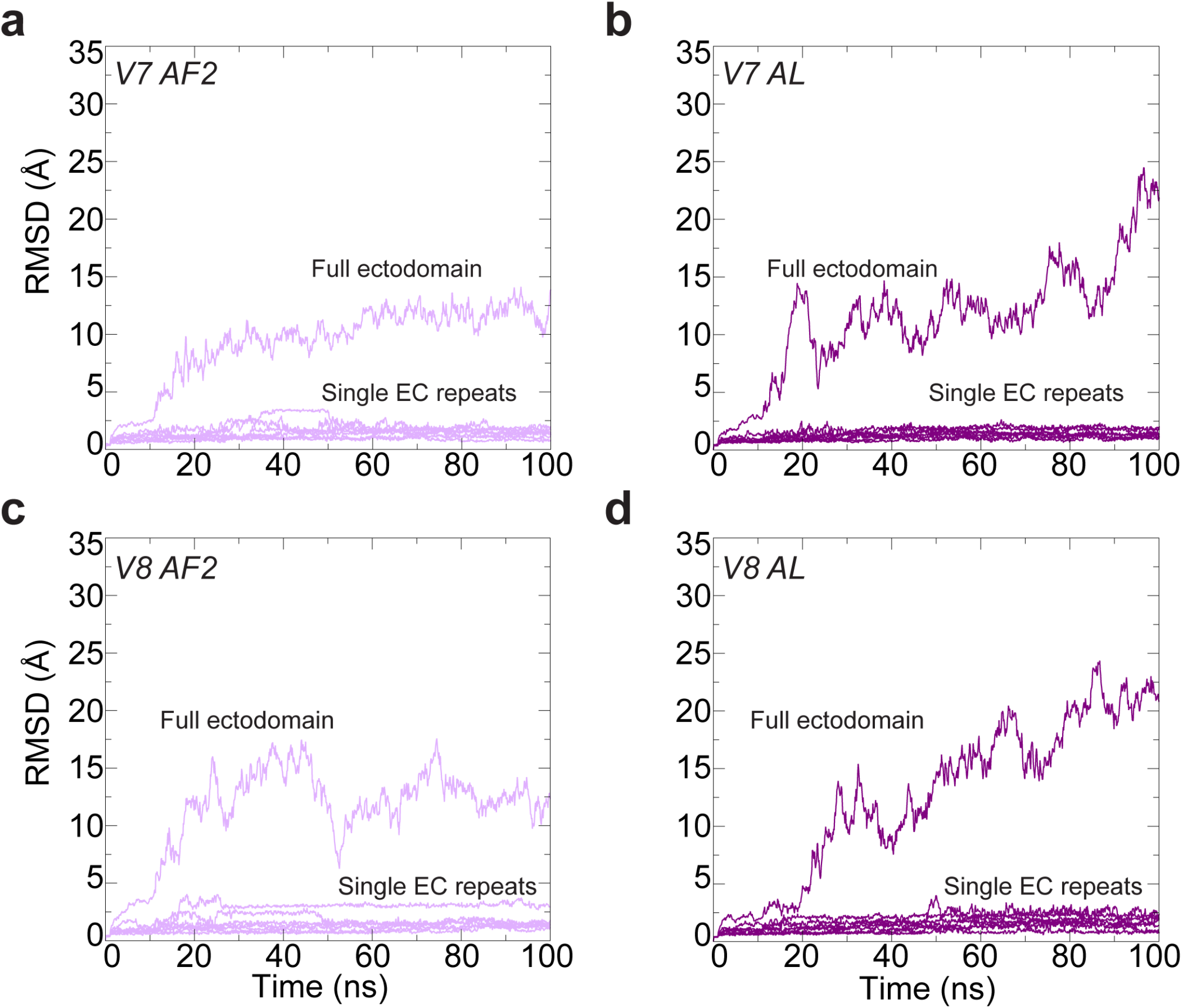
| Structural stability of mini-PCDH15s. a-d, Root-mean-square deviation (RMSD) as a function of time for the full ectodomains and individual EC repeats of the mini-PCDH15-V7 and -V8 AF2 and AL models.

**Fig. S3.**
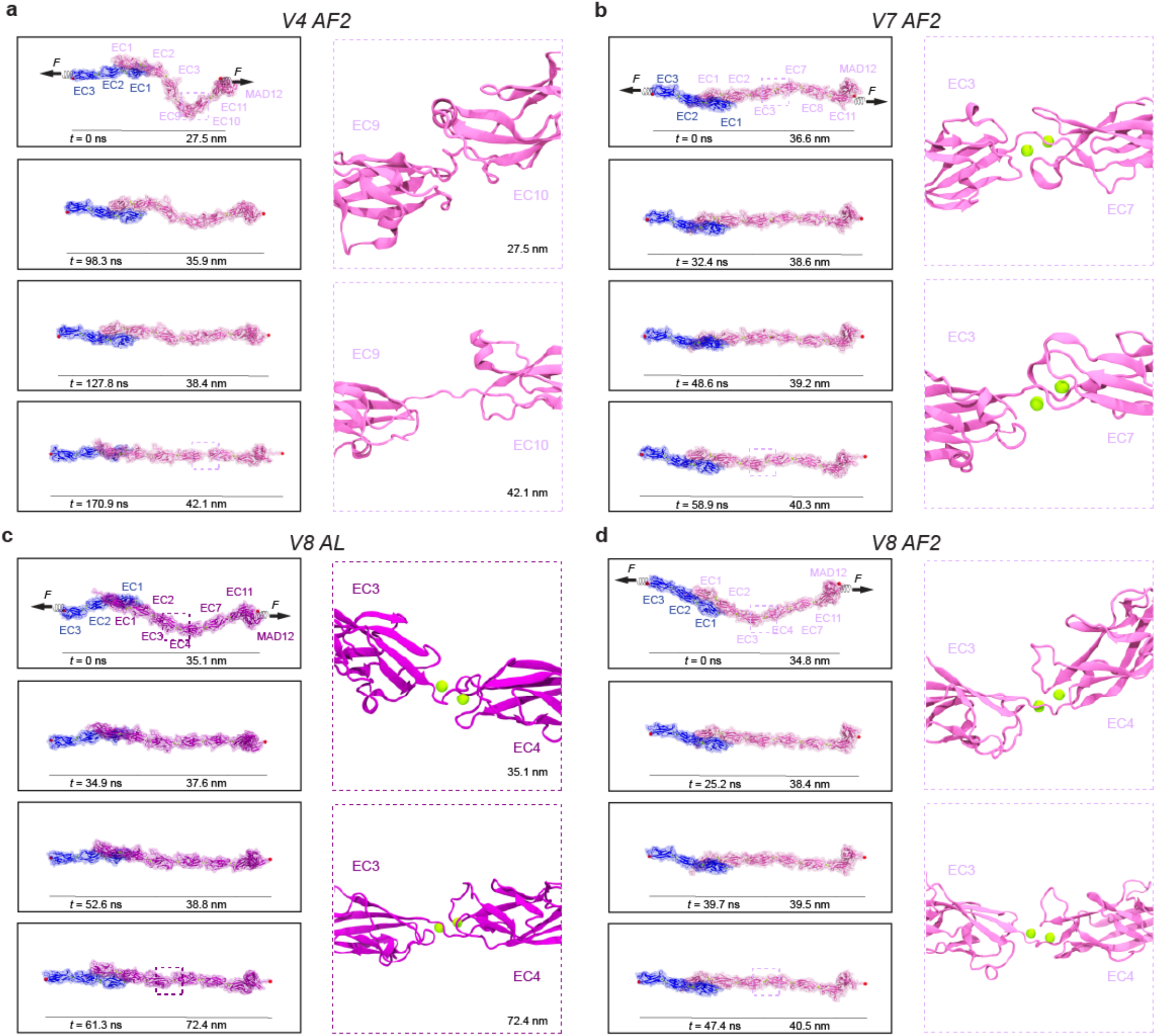
| **Elasticity of monomeric mini-PCDH15 proteins in complex with CDH23 EC1-3. a**, Snapshots of mini-PCDH15- V4 AF2 + CDH23 EC1-3 during stretching at 0.1 nm/ns (Sim1d). C-terminal C*α* atoms are shown as red spheres. Applied forces are indicated with spring arrows. Insets highlight the most extended EC linker (EC9-EC10). **b**, Snapshots of mini- PCDH15-V7 AF2 + CDH23 EC1-3 during stretching at 0.1 nm/ns (Sim3d). **c**, Snapshots of mini-PCDH15-V8 AL + CDH23 EC1-3 during stretching at 0.1 nm/ns (Sim6d). **d**, Snapshots of mini-PCDH15-V8 AF2 + CDH23 EC1-3 during stretching at 0.1 nm/ns (Sim5d).

**Fig. S4.**
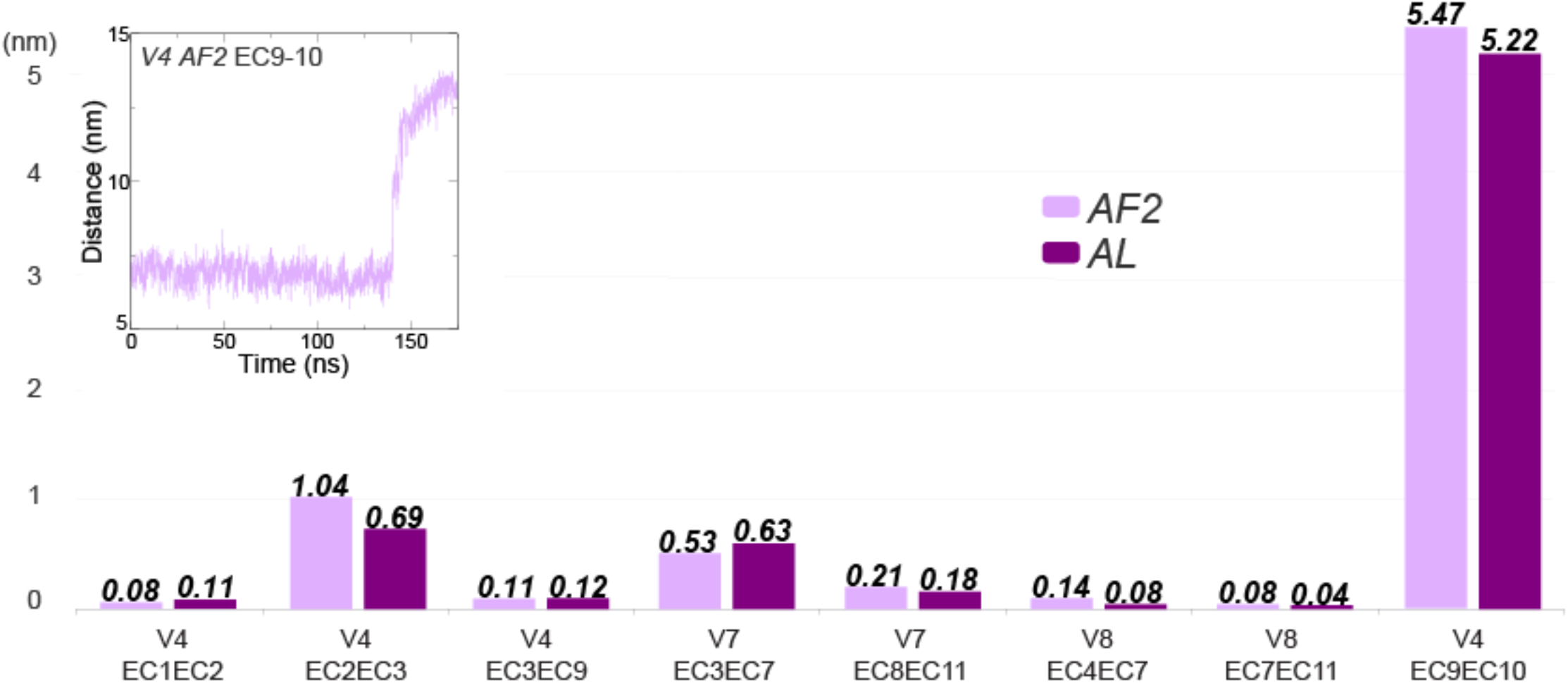
| Extension of mini-PCDH15 linker regions in stretching simulations at 0.1 nm/ns. The linker regions are depicted based on two different models: AF2 predictions shown in mauve and AL predictions in purple.

**Fig. S5.**
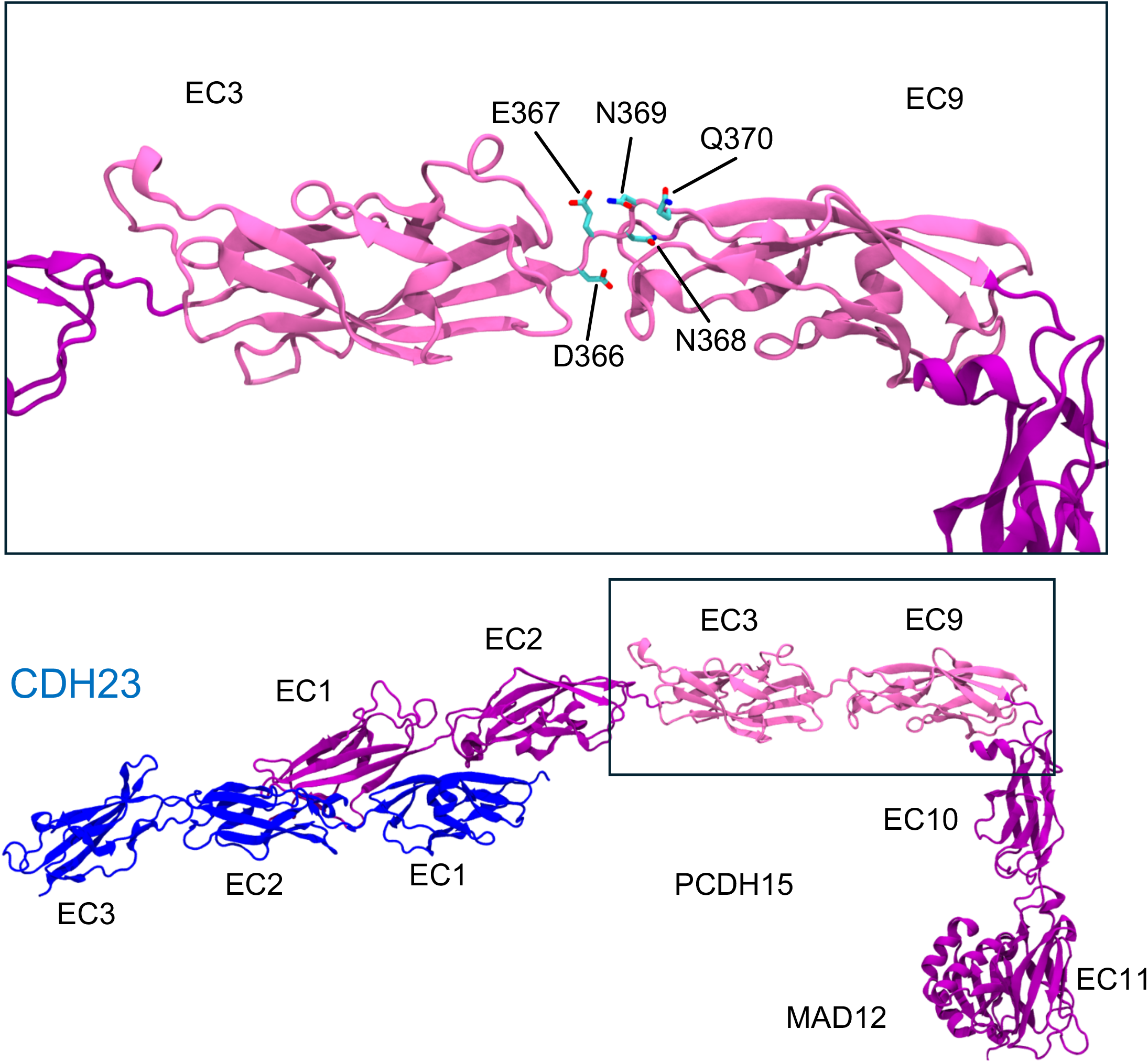
| **Assembly of mini-PCDH15-V4 AF2 model**. Base model from Choudhary et al. includes CDH23 EC1-3 in blue, PCDH15 EC1-2 in purple, and PCDH15 EC10-MAD12 in purple. Engineered linker EC3-EC9 (in mauve) was predicted using AF2. We then manually connected the C-terminal end of PCDH15 EC1-2 to the N-terminal end of the AF2-predicted PCDH15 EC3-EC9. Similarly, the C-terminal end of the AF2-predicted EC3-EC9 was connected to N-terminal end of PCDH15 EC10-MAD12. Box highlights the AF2-predicted EC3-EC9 subdomain.

**Fig. S6.**
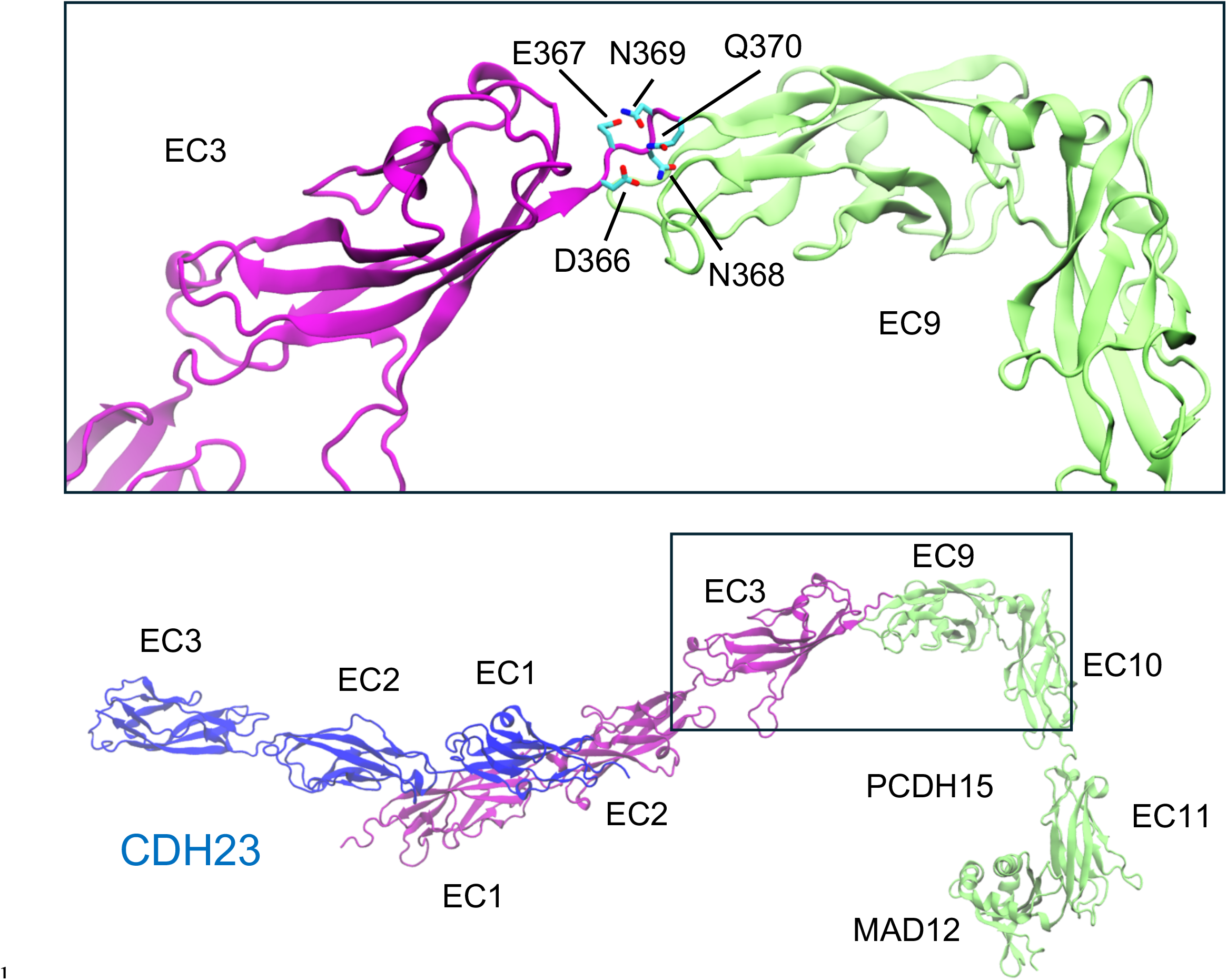
| **Assembly of mini-PCDH15-V4 AL model**. Base model from Choudhary et al. includes CDH23 EC1-3 in blue, PCDH15 EC1-3 in purple, and PCDH15 EC9-MAD12 in lime. We manually connected the C-terminal end of PCDH15 EC1- 3 to the N-terminal end of PCDH15 EC9-MAD12. Box highlights the engineered linker region.

**Table S1.** | RMSD values from COOT using atoms of protein backbone between PDB structures of linkers and WT domains (Å)

|  | 8TON EC3 | 8UMZ EC4<br>(chain A) | 8UMZ EC4<br>(chain B) | 8TON EC7 | 8UMZ EC7<br>(chain A) | 8UMZ EC7<br>(chain B) |
| --- | --- | --- | --- | --- | --- | --- |
| 5T4M EC3<br>(chain A) | 1.083 | - | - | - | - | - |
| 5T4M EC3<br>(chain B) | 0.842 | - | - | - | - | - |
| 5T4M EC4<br>(chain A) | - | 0.925 | 0.395 | - | - | - |
| 5T4M EC4<br>(chain B) | - | 0.395 | 0.473 | - | - | - |
| 5ULY EC3 | 0.511 | - | - | - | - | - |
| 5W1D EC4 | - | 0.729 | 0.719 | - | - | - |
| 5W1D EC7 | - | - | - | 0.516 | 0.594 | 0.649 |
| 6BWN EC7 | - | - | - | 0.399 | 0.435 | 0.533 |

**Table S2.** | Summary of simulations for mini-PCDH15 proteins + CDH23 EC1-3.

| Label | System | $t_{sim}$ (ns) | Type | Start | Speed (nm/ns) | Average force Peak (pN) | Size (#atoms) | Initial Sytem Size ($nm^3$) |
| --- | --- | --- | --- | --- | --- | --- | --- | --- |
| Sim1a | V4 AF2 | 100 | EQ | - | - | - | 1,239,195 | 76×14×12 |
| Sim1b |  | 1.24 | SMD | S1a | 10 | 695.0 +/- 197.8 |  |  |
| Sim1c |  | 9.85 | SMD | S1a | 1 | 388.4 +/- 23.0 |  |  |
| Sim1d |  | 170.9 | SMD | S1a | 0.1 | 355.8 +/- 12.7 |  |  |
| Sim2a | V4 AL | 100 | EQ | - | - | - | 938,545 | 76×16×8 |
| Sim2b |  | 1.2 | SMD | S2a | 10 | 713.5 +/- 261.7 |  |  |
| Sim2c |  | 9.77 | SMD | S2a | 1 | 390.7 +/- 51.2 |  |  |
| Sim2d |  | 189.2 | SMD | S2a | 0.1 | 315.1 +/- 0.6 |  |  |
| Sim3a | V7 AF2 | 100 | EQ | - | - | - | 1,060,405 | 76×12×12 |
| Sim3b |  | 0.94 | SMD | S3a | 10 | 754.4 +/- 236.0 |  |  |
| Sim3c |  | 6.74 | SMD | S3a | 1 | 431.9 +/- 49.0 |  |  |
| Sim3d |  | 84.0 | SMD | S3a | 0.1 | 291.8 +/- 48.8 |  |  |
| Sim4a | V7 AL | 100 | EQ | - | - | - | 1,060,492 | 76×12×12 |
| Sim4b |  | 0.98 | SMD | S4a | 10 | 729.5 +/- 236.2 |  |  |
| Sim4c |  | 6.35 | SMD | S4a | 1 | 411.9 +/- 55.9 |  |  |
| Sim4d |  | 100.6 | SMD | S4a | 0.1 | 287.2 +/- 4.0 |  |  |
| Sim5a | V8 AF2 | 100 | EQ | - | - | - | 1,060,357 | 76×12×12 |
| Sim5b |  | 0.96 | SMD | S5a | 10 | 753.8 +/- 204.0 |  |  |
| Sim5c |  | 6.16 | SMD | S5a | 1 | 402.9 +/- 65.7 |  |  |
| Sim5d |  | 84.5 | SMD | S5a | 0.1 | 353.3 +/- 41.2 |  |  |
| Sim6a | V8 AL | 100 | EQ | - | - | - | 1,060,501 | 76×12×12 |
| Sim6b |  | 0.96 | SMD | S6a | 10 | 731.3 +/- 204.8 |  |  |
| Sim6c |  | 6.25 | SMD | S6a | 1 | 388.7 +/- 68.2 |  |  |
| Sim6d |  | 84.1 | SMD | S6a | 0.1 | 309.7 +/- 7.7 |  |  |

**Table S3.** | Predicted elasticity of monomeric mini-PCDH15 proteins.

| Mini-PCDH15 Versions | Effective Spring Constant $k$ (mN/m) | | Extensibility (nm) | |
| --- | --- | --- | --- | --- |
| V4 AF2 | 3.3 | 34.2 | 9.0 | 5.0 |
| V4 AL | 3.3 | 17.2 | 7.0 | 7.5 |
| V7 AF2 | 36.3 |  | 4.5 |  |
| V7 AL | 33.0 |  | 5.5 |  |
| V8 AF2 | 53.9 |  | 3.5 |  |
| V7 AL | 55.5 |  | 4.0 |  |

**Table S4.** | RMSD values between AF2 and AF3 predictions for mini-PCDH15 engineered linkers.

| **Mini-PCDH15 Engineered linkers** | **RMSD Values Using Non-Hydrogen Atoms (Å)** | **RMSD Values Using Atoms of Protein Backbone (Å)** |
| --- | --- | --- |
| V4 AF2 EC3 | 2.359 | 1.166 |
| V4 AF2 EC3-EC9 linker | 2.555 | 0.698 |
| V4 AF2 EC9 | 1.773 | 0.441 |
| V7 AF2 EC3 | 2.236 | 0.943 |
| V7 AF2 EC3-EC7 linker | 1.779 | 0.083 |
| V7 AF2 EC7 | 1.815 | 0.189 |
| V7 AF2 EC7-EC8 linker | 1.662 | 0.069 |
| V7 AF2 EC8 | 1.779 | 0.314 |
| V7 AF2 EC8-EC11 linker | 1.489 | 0.071 |
| V7 AF2 EC11 | 2.661 | 1.005 |
| V8 AF2 EC4 | 1.721 | 0.192 |
| V8 AF2 EC4-EC7 linker | 1.798 | 0.103 |
| V8 AF2 EC7 | 1.752 | 0.122 |
| V8 AF2 EC7-EC11 linker | 1.649 | 0.091 |
| V8 AF2 EC11 | 3.221 | 1.714 |

## Notes

### Competing Interest Statement

The authors have declared no competing interest.

